# Modulation of RecFORQ- and RecA-mediated homologous recombination in *Escherichia coli* by isoforms of translation initiation factor IF2

**DOI:** 10.1101/2021.06.16.448654

**Authors:** Jillella Mallikarjun, L SaiSree, P Himabindu, K Anupama, Manjula Reddy, J Gowrishankar

## Abstract

Homologous recombination (HR) is critically important for chromosomal replication as well as DNA damage repair in all life forms. In *Escherichia coli*, the process of HR is comprised of (i) two parallel pre-synaptic pathways that are mediated, respectively, by proteins RecB/C/D and RecF/O/R/Q; (ii) a synaptic step mediated by RecA that leads to generation of Holliday junctions (HJs); and (iii) post-synaptic steps mediated sequentially by HJ-acting proteins RuvA/B/C followed by proteins PriA/B/C of replication restart. Combined loss of RuvA/B/C and a DNA helicase UvrD is synthetically lethal, which is attributed to toxicity caused by accumulated HJs since viability in these double mutant strains is restored by removal of the pre-synaptic or synaptic proteins RecF/O/R/Q or RecA, respectively. Here we show that, as in Δ*uvrD* strains, *ruv* mutations confer synthetic lethality in cells deficient for transcription termination factor Rho, and that loss of RecFORQ pre-synaptic pathway proteins or of RecA suppresses this lethality. Furthermore, loss of IF2-1 (which is one of three isoforms [IF2-1, IF2-2, and IF2-3] of the essential translation initiation factor IF2 that are synthesized from three in-frame initiation codons in *infB*) also suppressed *uvrD-ruv* and *rho-ruv* lethalities, whereas deficiency of IF2-2 and IF2-3 exacerbated the synthetic defects. Our results suggest that Rho deficiency is associated with an increased frequency of HR that is mediated by the RecFORQ pathway along with RecA. They also lend support to earlier reports that IF2 isoforms participate in DNA transactions, and we propose that they do so by modulation of HR functions.

**Importance:** The process of homologous recombination (HR) is important for maintenance of genome integrity in all cells. In *Escherichia coli*, the RecA protein is a critical participant in HR, which acts at a step common to and downstream of two HR pathways mediated by the RecBCD and RecFOR proteins, respectively. In this study, an isoform (IF2-1) of the translation initiation factor IF2 has been identified as a novel facilitator of RecA’s function in vivo during HR.

## Introduction

Genetically first identified as a mechanism to mediate exchange of hereditary determinants between cells (1), the process of homologous recombination (HR) in bacteria has also been recognized to play a crucially important role in maintenance of genomic integrity both during chromosomal replication and following DNA damage, just as is the case in archaea and eukaryotes [reviewed in (2–11)]. In *Escherichia coli*, protein RecA is central to the synaptic step of HR: RecA binds a suitable single-strand (ss)-DNA substrate to form a nucleoprotein filament that performs a homology search to enable annealing to a second suitable DNA molecule. This synaptic step is flanked by pre-synaptic and post-synaptic reactions, as described below.

The pre-synaptic reactions are designed to generate the ss-DNA substrate for RecA’s binding. Two alternative pre-synaptic pathways RecBCD and RecFOR (named after the principal proteins mediating them) exist to prepare substrates from, respectively, double-strand breaks (DSBs) and ss-gaps in DNA. Loss of either RecBCD or RecFOR pathway alone affects certain categories of DNA recombination and repair, whereas combined loss of both pathways confers a deficiency as severe as that with loss of RecA itself (3, 6).

The presence of RecA-bound nucleoprotein serves also as a trigger for an SOS response within the cell, by which several prophages are induced to enter lytic growth and genes of the LexA-repressed SOS regulon are transcriptionally activated. For the SOS response, RecA’s co-protease activity is stimulated following nucleoprotein assembly to facilitate auto-cleavage of LexA and prophage repressors [reviewed in (12)].

The post-synaptic reactions act to generate discrete recombinant DNA molecules following annealing of the DNA molecules and formation of Holliday junctions (HJs). The RuvAB helicase and RuvC resolvase are primary mediators at this step, which catalyze branch migration and resolution of HJs, respectively (3). HJ resolution can also be achieved in absence of RuvABC, in *E. coli* cells harboring a mutation (*rus-1*) that activates expression of the protein Rus encoded by a cryptic prophage (13, 14); however, the physiological role if any of Rus is unclear.

Included in the post-synaptic phase of HR are steps of replication restart, by which products generated after resolution of HJs are assimilated into the circular bacterial chromosome (or plasmid). Proteins mediating replication restart include PriABC, DnaT, and Rep, which are proposed to act through several alternative and redundant pathways (15–20).

Control of HR in *E. coli* is achieved by autoregulation (through the SOS response) (12), as well as by several other factors (3, 21). One of the latter is UvrD, which is a 3’-5’ DNA helicase that also serves as an anti-recombinase to disrupt RecA nucleoprotein filaments (22, 23). Loss of UvrD is associated with (i) a hyper-recombination phenotype mediated by the RecFORQ pre-synaptic pathway in HR assays, and (ii) synthetic lethality with *ruv* mutations, presumed to be the consequence of accumulation of trapped HJs leading to impedance of chromosomal replication (24–26). Both phenotypes are suppressed by loss of RecA, which also indicates that hyper-recombination is not essential for viability of *uvrD* mutants.

In this study we have identified that a deficiency of transcription termination factor Rho also confers synthetic lethality with *ruv* mutations and that, just as in the case with *uvrD-ruv* mutants, abrogation of the RecFORQ pathway or of RecA rescues this lethality. Rho is an essential protein in *E. coli* that mediates the termination of transcripts (other than rRNAs and tRNAs) which are not being simultaneously translated (27–31). Rho’s function has been implicated in the maintenance of genomic integrity (32), and in avoidance of RNA-DNA hybrids or R-loops from nascent untranslated transcripts including from antisense RNAs (27, 28, 30, 33, 34).

Another finding being reported from this study is that both *rho-ruv* and *uvrD-ruv* synthetic lethalities were suppressed by mutations in *infB*, that encodes the translation initiation factor IF2 (35); we also show that these *infB* mutants are down-regulated for HR functions. Studies of the past fifteen years from the Nakai group had already identified an intriguing connection between IF2 on the one hand and DNA transactions including DNA damage repair on the other (36–38). IF2 is essential for E. *coli* viability (35), and its function in translation initiation is conserved across bacteria, archaea, and eukaryotes (39, 40); a mammalian mitochondrial IF2 homolog can restore viability to an *E. coli* IF2 knockout strain (41). Three IF2 isoforms are synthesized in E. *coli* (see Supp. Fig. S3A), from in-frame initiation codons at positions 1,158, and 165 of the 890-codon-long *infB* ORF, that are herein designated as IF2-1, IF2-2 and IF2-3, respectively (42, 43). All isoforms are active for translation initiation and any one of them is sufficient for viability (44, 45). The Nakai lab had shown that isoforms IF2-1 and IF2-2,3 (the latter designation is used for the mixture of IF2-2 and IF2-3, since they are only marginally different from one another) behave differently in vitro in assays for phage Mu transposition, and that they confer differential tolerance in vivo to genotoxic agents. They had proposed that the isoforms exert varying influences on different pathways of replication restart (36–38).

In the accompanying paper (46), we show that *infB* mutants are also defective for two-ended DSB repair, which is mediated by RecA through the RecBCD-mediated pre-synaptic pathway. Based on these combined results, we propose that the IF2 isoforms differentially affect the efficiency of synapsis between DNA molecules during HR.

## Results

### The genetic assays to identify lethality of mutants and their suppression

The genetic (blue-white) assay to demonstrate lethality has been described earlier (27, 47–50). This method makes use of a single-copy-number-shelter plasmid encoding trimethoprim (Tp)-resistance and carrying *lacZ*^+^ as well as a functional copy of the gene(s) of interest, and whose partitioning into daughter cells during cell division is not stringently regulated. Consequently, when a strain with this plasmid, along with Δ*lac* and a mutation in the gene of interest on the chromosome, is grown in medium not supplemented with Tp, plasmid-free cells that arise spontaneously in the culture (at around 5 to 20%) will be able to grow as white colonies on Xgal-supplemented plates if and only if the now unsheltered chromosomal mutation is not lethal; on the other hand, control blue colonies (formed from cells retaining the shelter plasmid) would be observed as a majority in all situations. In the studies below, we have employed the blue-white assay with a shelter plasmid carrying either *rho*^+^ or both *rho*^+^ and *infB*^+^ genes to examine lethality or synthetic lethality of *rho*, *rho-ruv*, and *infB* genes.

### *rho-ruv* is synthetically lethal

As mentioned above, transcription termination factor Rho is essential for *E. coli* viability (27, 32). In this study, a *rho* mutant was obtained with an *opal* (TGA) chain-terminating missense substitution in codon 136, which was shown to be viable in strains carrying the *E. coli* K-12 version of the *prfB* gene encoding release factor 2 (RF2) but not in those carrying the *E. coli* B version (Supp. Fig. S1A i-ii); there is evidence that the former but not the latter permits a low frequency of stop codon-readthrough (51). This was validated by Western blot analysis, which showed, for the *rho-136^opal^* mutant, bands corresponding to both full-length (faint) and truncated Rho polypeptides (Supp. Fig. S1B). Another viable *opal* mutant in codon 157 of *rho* has been reported recently in *E. coli* K-12 (52).

The *rho-136^opal^* mutation was synthetically lethal with disruptions of the *ruv* genes (Δ*ruvA*, Δ*ruvC*, or Δ*ruvABC*) encoding the HJ enzymes RuvAB or RuvC, and the phenotype was manifested on both rich and defined media (see, for example, Fig. 1A ii-iii, Fig. 2i and Supp. Fig. S2 i). On the other hand, *rho*-136^*opal*^ was not lethal with *recA* (Fig. 1A iv). These results are consistent with those from an earlier report that *ruv*, but not *recA*, mutants are hyper-sensitive to the Rho inhibitor bicyclomycin (32).

**Figure 1:**
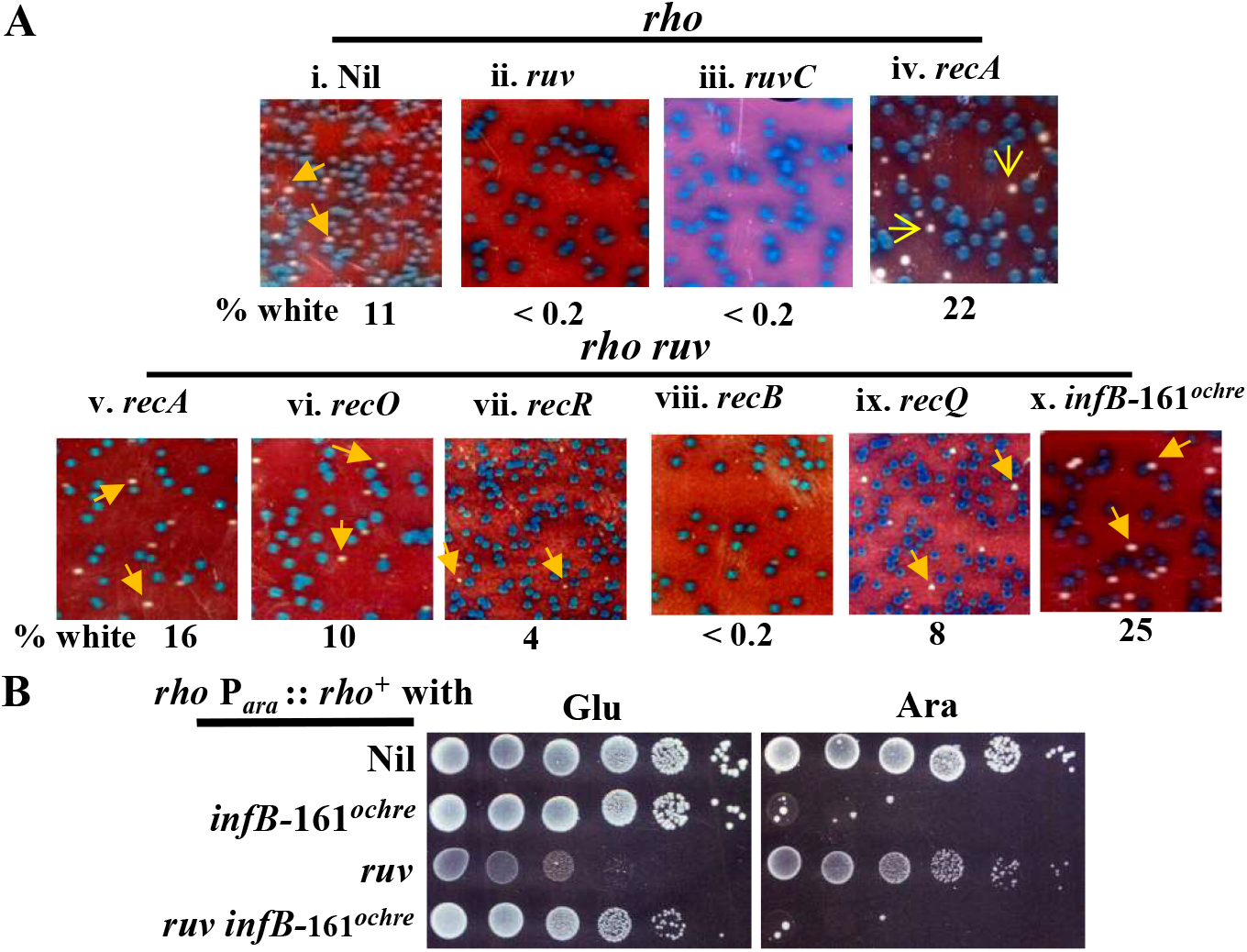
Synthetic lethality of *rho ruv* mutants, and their suppression by *infB*-161*^ochre^* or *rec* mutations. Mutant designations *rho* and *ruv* refer to alleles *rho*-136^*opal*^ and Δ*ruvABC*::Cm, respectively. All strain numbers mentioned are prefixed with GJ. (A) Blue-white screening assays, on defined medium at 30°, for strains carrying *rho*^+^ *infB*^+^ shelter plasmid pHYD5212. Representative images are shown; for each of the sub-panels, relevant chromosomal genotypes/features are given on top while the numbers beneath indicate the percentage of white colonies to total (minimum of 500 colonies counted). Examples of white colonies are marked by yellow arrows. Strains employed for the different sub-panels were pHYD5212 derivatives of: i, 15441; ii, 15447; iii, 19150; iv, 19845; v, 15460; vi, 15471; vii, 15491; viii, 15487; ix, 15497; and x, 15446. (B) Dilution-spotting assay, on minimal A supplemented with D-glucose (Glu) or Ara as indicated, of strains whose relevant genotypes/features are shown at left. Strains for different rows were (from top): 19379, 19380, 19381 (this strain was grown in Ara-supplemented medium before dilutions were spotted), and 19382.

**Figure 2:**
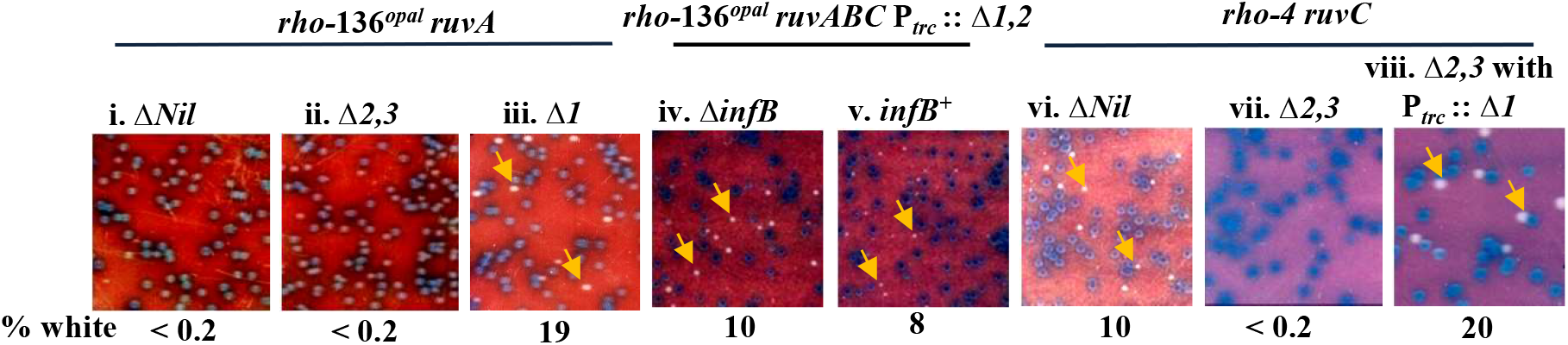
Modulation of *rho ruv* synthetic lethality by differential expression of IF2 isoforms. Blue-white screening assays were performed on defined medium at 30°, with strains carrying *rho*^+^ *infB*^+^ shelter plasmid pHYD5212. General notations are as described in legend to Figure 1A. All strains were Δ*infB*, with the exception of that for sub-panel v. Growth medium for sub-panels iv, v, and viii was supplemented with IPTG. Strains employed for the different sub-panels were pHYD5212 derivatives of (all strain numbers are prefixed with GJ): i, 15498; ii, 15500; iii, 15499; iv, 15458; v, 15457; vi, 19134; vii, 19131; and viii, 19132.

Synthetic *rho-ruv* lethality was suppressed by the *rus-1* mutation (Supp. Fig. S2 ii), which activates expression of the Rus resolvase from a cryptic prophage and thereby alleviates several *ruv* phenotypes (13, 14). The lethality was likewise suppressed by the *rpoB*35* mutation (53–55) (Supp. Fig. S2 iii), as also by ectopic expression of the (phage T4-encoded) R-loop helicase UvsW but not its active site mutant version UvsW-K141R (56, 57) (Supp. Fig. S2 iv-v). Both *rpoB*35* and UvsW alleviate the deleterious effects of Rho deficiency and can rescue Δ*rho* lethality (27, 28, 32).

### Loss of RecA, or of RecFOR pathway components, suppress *rho-ruv* lethality

Given the observations that the *rho-136^opal^* mutation was lethal with *ruv* but not with *recA*, the possibility was considered that excessive (and unnecessary) HR triggered in presence of the *rho* mutation was leading to accumulation of HJ intermediates in absence of RuvABC, resulting in cell death. Indeed, we could show that *rho-ruv* lethality is also rescued by Δ*recA* (Fig. 1A v, and Supp. Fig. S2 vii). An alternative explanation, that death was on account of excessive RecA-dependent SOS induction in *rho-ruv* mutants, was excluded since lethality was not rescued by the *lexA3* mutation [which encodes a LexA variant that is non-cleavable by RecA, and hence also abolishes SOS induction (12)] (Supp. Fig. S2 viii).

We then tested which of the two pre-synaptic pathways putatively feeds into the process of excessive HR in *rho-ruv* mutants. The synthetic lethality was suppressed by mutations in *recO* or *recR* (of the RecFOR pathway) (Fig. 1A vi-vii), but not by mutation in *recB* (of the RecBC pathway) (Fig. 1A viii). RecQ helicase activity is implicated in RecFOR pathway function (3, 6), and Δ*recQ* also was a suppressor of *rho-ruv* lethality (Fig. 1A ix).

### Suppression of *rho-ruv* lethality by loss of IF2-1

A new suppressor of *rho-ruv* lethality on both defined and rich media (Fig. 1A x, and Supp. Fig. S2 vi) was obtained and characterized. It was mapped to the *infB-nusA* locus, and was shown by DNA sequencing to be an *ochre* (TAA) nonsense codon mutation at position 161 of the *infB* ORF. One would expect, from its location in the *infB* ORF, that this mutation abrogates synthesis of the IF2-1 and IF2-2 isoforms, which was confirmed by Western blotting (Supp. Fig. S3C, lane 6).

By itself, the *infB-161^ochre^* mutation was lethal (Supp. Fig. S3B i) but it could be rescued by *rho* mutation (Supp. Fig. S3B ii). We interpret these findings as indicative of the notions (i) that nonsense substitution at codon 161 in *infB* induces Rho-mediated premature transcription termination within the gene, and therefore (ii) that synthesis of the lone IF2-3 isoform to ensure viability is itself contingent on relief of transcriptional polarity conferred by *rho* mutation.

Thus, viability of a triple mutant *rho ruv infB-161^ochre^* is based on mutual suppression (i) of *infB-*161^*ochre*^ lethality by *rho*, and (ii) of *rho ruv* lethality by *infB-161^ochre^.* This inference was supported by findings from experiments in which expression of an ectopically located *rho^+^* gene was placed under control of an L-arabinose (Ara)-inducible promoter (47) (Fig. 1B): that a *rho-ruv* derivative (row 3) is viable only on Ara-supplemented medium, whereas an *infB-161^ochre^-rho* mutant (row 2) as well as the triple mutant *infB-161^ochre^-rho-ruv* (row 4) are viable only on medium not supplemented with Ara. The *infB-rho* mutant was inhibited for growth at 22° (Supp. Fig. S3D), consistent with the known requirement for IF2 at low temperatures (58).

### Suppression of *rho-ruv* lethality by ectopic *infB* constructs

Based on the suppressor characterization results above, we surmised that it is the loss of isoforms IF2-1 or IF2-2 (or both together), which confers suppression of *rho-ruv* lethality. Accordingly, we then tested two other sets of ectopic chromosomal *infB* constructs for their ability to rescue *rho-ruv* lethality, in derivatives carrying a deletion of the native *infB* locus. (In the descriptions below, the designations *infB^+^* and Δ*infB* are used for the wild-type and deletion alleles, respectively, at the native chromosomal location).

In one set of three ectopic constructs, which has been described earlier by Nakai and coworkers (37, 38), IF2 expression remains under control of the natural *cis* regulatory elements for *infB* but not all isoforms are expressed from the different constructs (designation used in this study for each of them given in parentheses): that encodes only IF2-1, but not IF2-2 or IF2-3 (Δ*2*,*3*); that encodes both IF2-2 and IF2-3, but not IF2-1 (Δ*1*); and that encodes all three isoforms (Δ*Nil*). The constructs were validated for expression of the different IF2 isoforms by Western blot analysis (Supp. Fig. S3C, lanes 2 to 5 and lane 9). Upon testing in the *rho-ruv* strains, our results showed that the Δ*1* construct, but not Δ*2,3* or Δ*Nil*, could suppress *rho-ruv* synthetic lethality (Fig. 2 and Supp. Fig. S4, compare in each case iii with i and ii). These findings are consistent with that of *rho-ruv* suppression by *infB-161^ochre^*, since the latter also fails to express isoform IF2-1.

The second set of ectopic chromosomal constructs were prepared in this work and were designed for differential expression of IF2 isoforms from an isopropyl-β-D-thiogalactoside (IPTG)-inducible heterologous P_trc_ promoter (designation used for each given in parentheses): that expresses both IF2-2 and IF2-3, but not IF2-1 (P_*trc*_-Δ*1*); and that expresses IF2-3 alone (P_*trc*_-Δ*1*,*2*) (see Supp. Fig. S3C, lanes 8 and 7, respectively, for Western blot confirmation). Both constructs could suppress *rho-ruv* lethality (Fig. 2 iv and Supp. Fig. S4 iv-v), indicating once again that lethality suppression occurs when isoform IF2-1 is absent from the cells. Similar results were obtained irrespective of which *ruv* mutation (Δ*ruvA*, Δ*ruvC*, or Δ*ruvABC*) was used in the experiments (Fig. 2 iv; and Supp. Fig. S4 iv-v). With induced IF2-3 expression from the IPTG-regulated construct, suppression was obtained even in derivatives carrying the native *infB^+^* locus, but the colony growth was less robust (Fig. 2 v).

These results indicate that it is an imbalance between levels of isoforms IF2-1 (low) and IF2-2,3 (high) that determines suppression of *rho-ruv* lethality. This suppression is not because of a restoration (or bypass) of Rho or Ruv function in these strains, since loss of IF2-1 did not reverse (i) the phenotypes associated with *rho* mutation (33), such as a Gal^+^ phenotype that follows relief of premature transcription termination in the *gal* operon caused by *galEp3* mutation (Supp. Fig. S1C, compare iii-iv with ii), or lethality caused by runaway replication of plasmid pACYC184 (Supp. Fig. S1A iii-v); nor (ii) the UV-sensitivity known for *ruv* mutants (3, 59), and indeed it aggravated the latter phenotype (Supp. Fig. S5A).

### Loss of IF2-2,3 exacerbates *rho-ruv* defect

The following experiments lent additional support to the notion that *rho-ruv* lethality is modulated by the balance between isoforms IF2-1 and IF2-2,3. Derivatives with the missense *rho-4* mutation (encoding Rho-A243E (33), see Supp. Fig. S1B lane 2 for Western blot) may be expected to be less compromised for Rho function than those with *rho*-136^*opal*^, and a test for intracellular R-loop prevalence with monoclonal antibody S9.6 (28, 60) showed this to be so (Supp. Fig. S1D). Unlike *rho*-136^*opal*^, the *rho-4* mutation was not synthetically lethal with *ruv* in the Nakai Δ*Nil* strain expressing all three IF2 isoforms (Fig. 2 vi). However, this combination (*rho-4 ruv*) conferred lethality in the Δ*2*,*3* derivative (lacking IF2-2,3) (Fig. 2 vii), and this lethality was rescued by IPTG-induced expression of IF2-2,3 (Fig. 2 viii). Thus, absence of IF2-1 and of IF2-2,3 exert the apparently opposite phenotypes of, respectively, alleviating and exacerbating the RecFORQ- and RecA-mediated sickness of *rho-ruv* mutants.

### Synthetic lethality of *uvrD-ruv* is also suppressed by loss of IF2-1

Several groups have earlier reported synthetic *uvrD-ruv* lethality (24–26), which is suppressed by *recA* and *recFORQ* but not *recBC* (that is, very similar to our findings with *rho-ruv* lethality). The mechanism invoked has been that of excessive and unnecessary HR in the *uvrD* mutant, leading to accumulation of toxic recombination intermediates in absence of RuvABC.

To test whether IF2 isoforms affect *uvrD-ruv* lethality, we expressed (from a doxycycline [Dox]-inducible promoter) a dominant-negative RuvC protein designated as RDG, that binds and traps HJs (59); these experiments were done in a set of Δ*uvrD* Δ*infB* derivatives each carrying one of the ectopically integrated Nakai constructs for different IF2 isoforms. Viability of the Δ*Nil* and Δ*2*,*3* strain derivatives was reduced 10^3^- and 10^4^-fold, respectively, upon Dox addition whereas the Δ*1* derivative was only minimally affected (Fig. 3). A *recA* derivative of the Δ*Nil* strain survived Dox, as too did the control *uvrD*^+^ *recA*^+^ strain (Fig. 3, last and first rows, respectively). These results confirm that *uvrD-ruv* is synthetically lethal (more severely so in absence of IF2-2,3), and that this lethality is rescued upon loss of RecA or of IF2-1.

**Figure 3:**
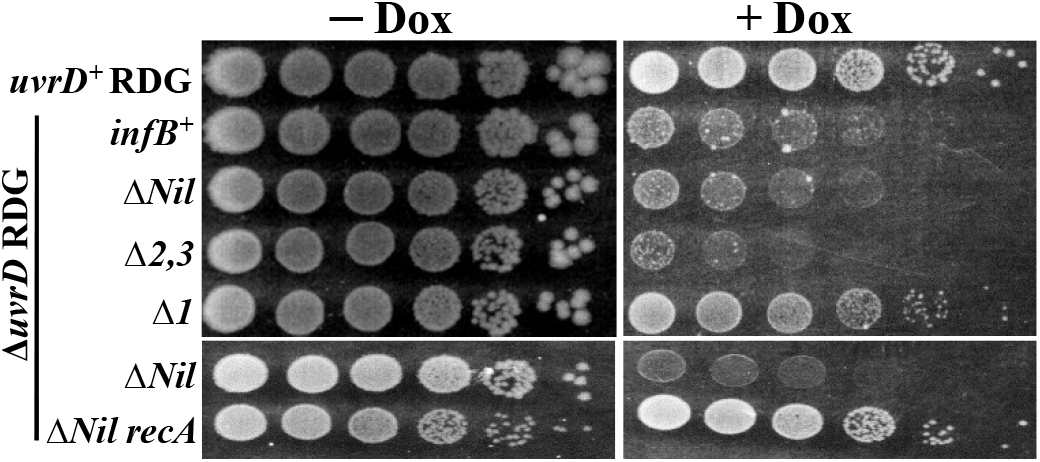
Synthetic lethality conferred by dominant negative *ruvC* mutation (RDG) in Δ*uvrD* mutant, and its suppression by loss of IF2-1. Dilution-spotting assay was performed on LB medium without (−) and with (+) Dox supplementation, of RDG-bearing derivatives whose relevant genotypes/features are indicated at left; strains on all but the top two rows were also Δ*infB*. Strains employed for different rows were (from top, all strain numbers are prefixed with GJ): 19127, 19161, 19801, 19802, 19803, 19843, and 19835.

### HR frequency is oppositely affected by loss of IF2-1 and of IF2-2,3, and is elevated in *rho* mutants

The results above had indicated that loss of IF2-1 phenocopies the loss of RecA or of the pre-synaptic RecFORQ pathway to confer suppression of *rho-ruv* and *uvrD-ruv* lethalities. We then tested the differential effects, if any, of IF2 isoforms on recovery of recombinants following HR, for which we employed several assays such as those of phage P1 transduction, inter-plasmidic recombination (that leads to reconstitution of a tetracycline (Tet)-resistance gene from two partially overlapping deletion alleles) (61–64), and the Konrad assay (that similarly entails reconstitution of an intact *lacZ* gene from a split pair of partially overlapping *lacZ* fragments located at distant sites on the chromosome) (65, 66). Recombination events in each of the assays above are RecA-dependent (3, 61, 65); inter-plasmidic recombination is mediated by the RecFOR pre-synaptic pathway (61, 63, 64), whereas P1 transduction and recombination in the Konrad assay are RecBCD-dependent (65).

In the HR assays, loss of IF2-1 and of IF2-2,3 were associated with moderate reduction and elevation, respectively, in recovery of recombinants in comparison with values for the Δ*Nil* strain (Fig. 4A-C). A moderate reduction in P1 transduction frequency for the strain lacking IF2-1 has also been reported earlier (37).

**Figure 4:**
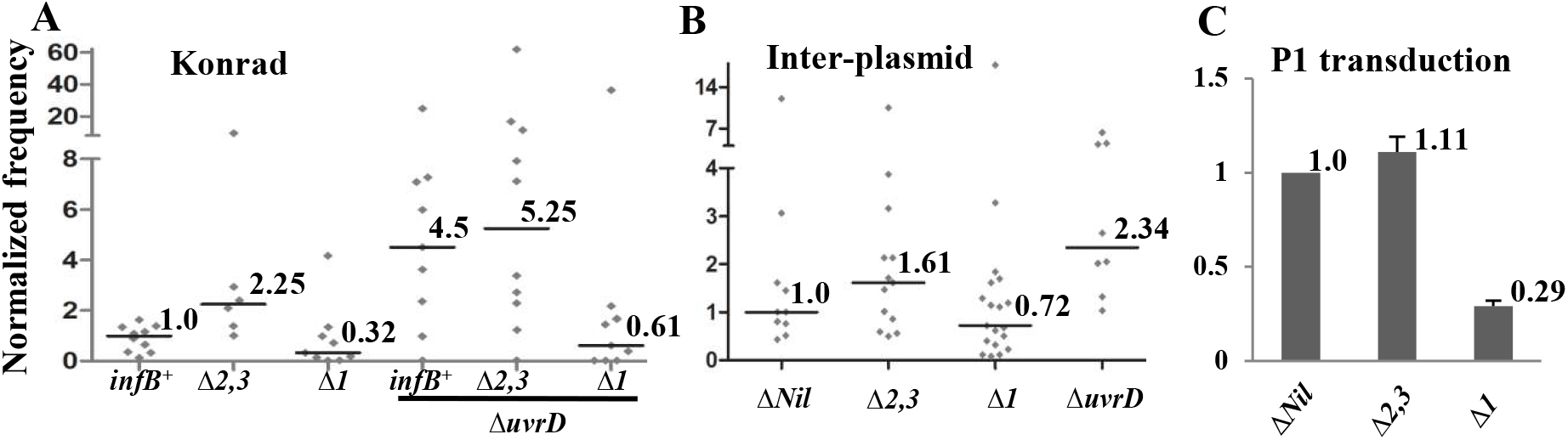
Effect of IF2 isoforms on HR. Recombination frequency data are given for strains with the indicated genotypes / features in the Konrad (A), inter-plasmid recombination (B), and P1 transduction (C) assays, after normalization to the value for the cognate control strain (taken as 1, and shown at extreme left for each panel); the actual control strain values were 2.8*10^−7^/viable cell, 6*10^−4^/viable cell, and 1.5*10^−5^/phage, respectively. In panels A and B, values from every individual experiment is shown, and median values are given beside the denoted horizontal lines. Panel C depicts the mean value (given alongside each bar) and standard error for each strain. In all panels, strains whose designations include Δ*Nil*, Δ1, or Δ*2*,*3* were also Δ*infB*. Strains used were (from left, all strain numbers mentioned are prefixed with GJ unless otherwise indicated): in panel A, SK707, 19171, 19186, 19162, 19184, and 19165; in panel B, 19197, 19196, 19195, and 19847; and in panel C, 19193, 19194, and 15494.

As expected (22, 65, 66), loss of UvrD conferred a hyper-recombination phenotype (Fig. 4A-B), but the HR frequency in a derivative that had lost both UvrD and IF2-1 was once again low and resembled that in a strain lacking IF2-1 alone (Fig. 4A, compare columns 3 and 6). Thus, loss of IF2-1 is epistatic to Δ*uvrD*, indicating that UvrD’s function in HR precedes and is in the same pathway as that of IF2-1, but that the two act oppositely.

Since the results above suggested that loss of IF2-1 is associated with reduced RecA function in HR (see *Discussion*), we examined whether there is reduction in SOS induction [which is mediated by RecA’s coprotease activity that is activated upon binding to ss-DNA (12)] as well in this situation. The results indicate that, as measured by *sulA-lac* expression (67), the SOS response (both basal as well as that following DNA damage with phleomycin) is not decreased and may in fact be modestly elevated in the Δ*1* strain (Supp. Fig. S5B).

Our finding of *rho-ruv* synthetic lethality that is suppressed by *recA* also suggests, based on its parallels with *uvrD-ruv* lethality, that non-essential HR occurs at elevated frequency in the *rho-136^opal^* mutant. Measurements of HR frequency, both by the Konrad assay and by conjugation, indicate that the *rho* mutant does exhibit a moderate increase in HR frequency (Supp. Fig. S1E).

## Discussion

The major findings of this study are (i) that Rho deficiency is synthetically lethal with *ruv* mutations in a manner that is similar to *uvrD-ruv* lethality (with trapped HJs, generated through the RecFORQ pre-synaptic pathway and RecA, being responsible for inviability in both cases); and (ii) that an imbalance of isoforms of the translation initiation factor IF2 affects HR functions in the cells. Each of these is further discussed below.

### Why is *rho-ruv* lethal?

Several features are shared between synthetic lethalities *rho-ruv* and *uvrD-ruv*, suggesting commonality of mechanisms in the two instances. Thus, both lethalities require proteins RecA, RecFORQ, and translation initiation factor isoform IF2-1.

In case of *uvrD-ruv*, the model proposed by Rosenberg and colleagues (24) is that in absence of UvrD, there is excessive but unnecessary HR through the RecFORQ pathway, which then renders RuvABC essential for resolving the ensuing HJ intermediates. Likewise for the *rho* mutant, we suggest that on account of an increased prevalence of R-loops with displaced ss-DNA, increased HR is triggered through the RecFOR pathway thus necessitating RuvABC’s presence for viability (Fig. 5, panels a-c).

**Figure 5:**
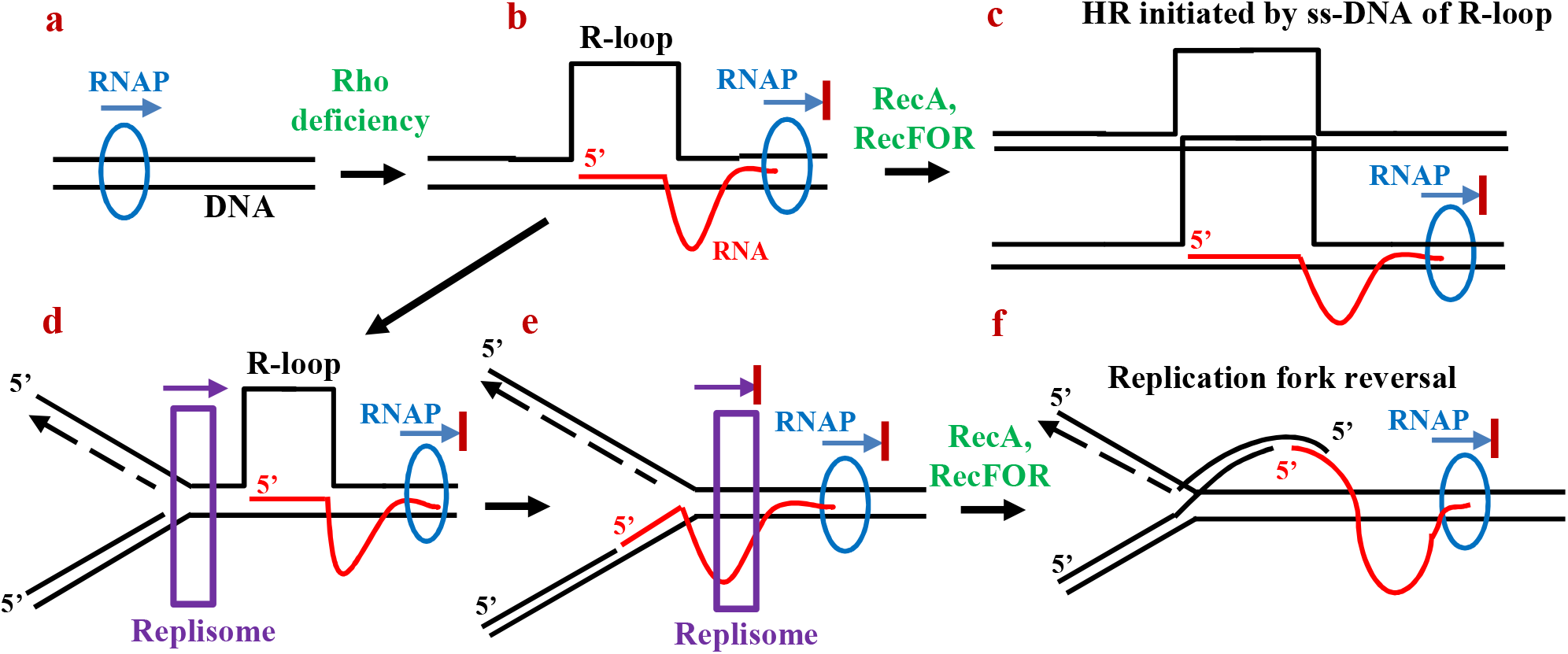
Two models to explain HJ formation (and hence need for RuvABC) in *rho* mutants. **(a-b)** Rho deficiency provokes R-loop generation from nascent untranslated transcripts, associated with RNA polymerase (RNAP) backtracking and arrest. **(c)** HR is initiated by invasion of ss-DNA of R-loop into a sister chromosome, mediated by RecFOR and RecA. **(d-e)** Co-directional replisome traverses R-looped region before it is arrested by backtracked RNAP. **(f)** Replisome disassembly followed by RecA- and RecFOR-mediated replication fork reversal generates HJ, with the RNA-DNA hybrid now as part of the extruded tail.

An alternative model for *rho-ruv* lethality, that also invokes R-loops, is that depicted in Figure 5, panels d to f. In this model, a chromosomal R-loop in the *rho* mutant serves to inhibit replisome progression (68), which is then followed by RecA- and RecFOR-mediated replication fork reversal (69–71). Since the extruded duplex includes an RNA-DNA hybrid, the fork cannot be restored through RecBCD action and hence the action of RuvABC at the HJ becomes essential for viability. The two models are not mutually exclusive.

In both models, UvsW expression and *rpoB*35* are suppressors of *rho-ruv* lethality because they presumably act to reduce R-loop prevalence in Rho-deficient strains (27, 28, 32).

### Opposing effects of IF2-1 and IF2-2,3 isoforms in HR pathways

Early studies had established that isoforms IF2-1 and IF2-2,3 are together required for optimal growth of *E. coli* (45). Nakai and coworkers (37, 38) have reported that loss of IF2-1 or of IF2-2,3 is each associated with sensitivity to different kinds of DNA damage.

In our study, loss of IF2-1 or of IF2-2,3 was associated with suppression or aggravation, respectively, of *rho-ruv* and *uvrD-ruv* sickness. The model for opposing effects of these isoforms is supported also by our findings in HR assays. At a mechanistic level, however, it is unclear whether a particular phenotype is caused by absence of one IF2 isoform or exclusive presence of another.

### How do IF2 isoforms influence HR?

The studies reported in this and in the accompanying paper (46), taken together, have identified IF2 isoforms as novel players in HR, but the mechanisms by which they act for this purpose are unknown. The present study has shown that loss of IF2-1 (i) phenocopies loss of the RecFORQ pre-synaptic pathway and of RecA in suppressing *rho-ruv* and *uvrD-ruv* lethalities, and (ii) reduces the recovery of recombinants following HR. In the accompanying paper (46), it is shown that loss of IF2-1 confers profound sensitivity to two-ended DSBs, whose repair is mediated by the RecBCD pre-synaptic pathway along with RecA.

The model we propose is that absence of IF2-1 leads directly or indirectly to reduction in efficiency of a step in HR which is (i) downstream of (and common to) the RecBCD and RecFOR pathways, and (ii) upstream of the post-synaptic reactions mediated by RuvABC. Formation of the RecA nucleoprotein itself is apparently unaffected, since the SOS response (12) is not perturbed by loss of IF2-1. In a further elaboration of this model in the accompanying paper (46), it is the strength of RecA-mediated annealing between a pair of homologous DNA molecules that is postulated to be decreased in absence of IF2-1.

The proposed model is consistent with our finding that IF2-1’s role in HR is downstream to that of UvrD, which acts to modulate efficiency of RecA nucleoprotein formation on DNA substrates generated by the two alternative pre-synaptic pathways (22, 23). Deficiency of IF2-2,3 is proposed to have the opposite effect, of enhancing the efficiency of the same step in HR as that diminished by loss of IF2-1.

Our model that loss of IF2-1 compromises RecA function is different from that of Nakai and colleagues (37, 38), who proposed that IF2 isoforms differentially influence different replication restart pathways (which are post-synaptic, and downstream of RuvABC action). As explained above, our finding that loss of IF2-1 phenocopies the *recA* mutation in suppressing *rho-ruv* and *uvrD-ruv* lethalities can best be explained only by invoking a role for IF2-1 prior to the step of RuvABC action.

To test the models above, in vitro studies may be needed to examine whether IF2 isoforms act directly to modulate HR, and to determine their precise role(s) in the process. Regulation of HR and of recombinational repair functions is important in both prokaryotes and eukaryotes (10, 72), and factors previously identified for such regulation in *E. coli* include UvrD, mismatch repair proteins, DinI, and RecX (3, 21, 22, 73, 74).

## Materials and Methods

### Growth media, bacterial strains and plasmids

The routine rich and defined growth media were, respectively, LB and minimal A with 0.2% glucose (75) and, unless otherwise indicated, the growth temperature was 37°. Supplementation with Xgal and with antibiotics ampicillin (Amp), chloramphenicol (Cm), kanamycin (Kan), spectinomycin (Sp), Tet, and Tp were at the concentrations described earlier (47). Phleomycin supplementation was at 3 μg/ml. For induction of gene expression from the appropriate regulated promoters, Ara, Dox, and IPTG were added at 0.2%, 50 ng/ml, and 0.5 mM, respectively. *E. coli* strains used are listed in Supplementary Table S1, with the following knockout (Kan^R^ insertion-deletion) alleles sourced from the collection of Baba et al. (76): *dinF*, *ilvA*, *leuA*, *leuD*, *racC*, *recA*, *recB*, *recO*, *recQ*, *recR*, *ruvA*, *serA*, *thrA*, *uvrD*, *ybfP*, and *yihF*; the Δ*infB* knockout mutation has also been described earlier (41).

Plasmids described earlier include pBR322 (Tet^R^ Amp^R^, ColE1 replicon) (77); pACYC184 (Tet^R^ Cm^R^, p15A replicon) (78); pCL1920 (Sp^R^, pSC101 replicon) (79); pMU575 (Tp^R^, single-copy-number vector with *lacZ^+^*) (80); pHYD2411 (Tp^R^, pMU575 derivative with *rho^+^*) (27); and pTrc99A (Amp^R^, for IPTG-inducible expression of gene of interest) (81). Plasmids pKD13 (Kan^R^ Amp^R^), pKD46 (Amp^R^), and pCP20 (Cm^R^ Amp^R^), for use in recombineering experiments and for Flp-mediated site-specific excision of FRT-flanked DNA segments, have been described by Datsenko and Wanner (82). Plasmids constructed in this study are described in the *Supplementary Text*.

### Methods

Procedures for P1 transduction (83), recombineering on the chromosome or plasmids (82), determination of UV tolerance (75), and R-loop detection with S9.6 monoclonal antibody (28) were as described. The Miller protocol was followed for β-galactosidase assays (75), and enzyme specific activity values are reported in the units defined therein. Protocols of Sambrook and Russell (84) were followed for recombinant DNA manipulations, PCR, and transformation. The Western blotting procedure, with rabbit polyclonal anti-IF2 or anti-Rho antisera (kind gifts from Umesh Varshney and Ranjan Sen, respectively), was essentially as described (49). Chromosomal integration, at the phage λ *att* site, of pTrc99A derivatives expressing IF2-2,3 or IF2-3, was achieved by the method of Boyd et al. (85). HR assay methods are described in the *Supplementary Text*, and include those based on conjugation (75), the Konrad assay (65), and inter-plasmid recombination (61, 62).

## Supporting information

Supplementary Data including Supplementary Text, Supplementary References, Supplementary Table S1, and Supplementary Figures S1-S5.

## Supplemental material

Supplemental material is provided as a PDF file “Supplemental File 1”.

## Acknowledgements

We thank R Harinarayanan, Hiroshi Nakai, Susan Rosenberg, Ranjan Sen, and Umesh Varshney for strains, plasmids, and reagents; Anjana Badrinarayanan, Rachna Chaba, Dipak Dutta, and Mohan Joshi for comments on the manuscript; and COE team members for advice and discussions.

This work was supported by Government of India funds from (i) DBT Centre of Excellence (COE) project for Microbial Biology – Phase 2, (ii) SERB project CRG/ 2018/ 000348, and (iii) DBT project BT/ PR34340/ BRB/ 10/ 1815/ 2019. JM was recipient of a DST-INSPIRE fellowship, and JG was recipient of the J C Bose fellowship and INSA Senior Scientist award.

We declare that there are no conflicts of interest.

## Notes

### Competing Interest Statement

The authors have declared no competing interest.

### Summary of Updates

This is a companion article to the article titled as "Essential role for an isoform of Escherichia coli translation initiation factor IF2 in repair of two-ended DNA double-strand breaks"

## References

1. Clark AJ. 1973. Recombination deficient mutants of *E. coli* and other bacteria. Annu Rev Genet 7:67–86.

2. Kuzminov A. 1995. Collapse and repair of replication forks in *Escherichia coli*. Mol Microbiol 16:373–384.

3. Kuzminov A. 1999. Recombinational Repair of DNA Damage in *Escherichia coli* and Bacteriophage λ. Microbiol Mol Biol Rev 63:751–813.

4. Cox MM, Goodman MF, Kreuzer KN, Sherratt DJ, Sandler SJ, Marians KJ. 2000. The importance of repairing stalled replication forks. Nature 404:37–41.

5. Sinha AK, Possoz C, Leach DRF. 2020. The Roles of Bacterial DNA Double-Strand Break Repair Proteins in Chromosomal DNA Replication. FEMS Microbiol Rev 44:351–368.

6. Kowalczykowski SC. 2000. Initiation of genetic recombination and recombination-dependent replication. Trends Biochem Sci 25:156–165.

7. Lenhart JS, Schroeder JW, Walsh BW, Simmons LA. 2012. DNA Repair and Genome Maintenance in *Bacillus subtilis*. Microbiol Mol Biol Rev 76:530–564.

8. Costes A, Lambert SAE. 2013. Homologous recombination as a replication fork escort: Fork-protection and recovery. Biomolecules 3:39–71.

9. Bianco PR, Lu Y. 2021. Single-molecule insight into stalled replication fork rescue in *Escherichia coli*. Nucleic Acids Res 49:4220–4238.

10. Kowalczykowski SC. 2015. An Overview of the Molecular Mechanisms of Recombinational DNA Repair. Cold Spring Harb Perspect Biol 7:a016410.

11. Bell JC, Kowalczykowski SC. 2016. Mechanics and Single-Molecule Interrogation of DNA Recombination. Annu Rev Biochem 85:193–226.

12. Simmons LA, Foti JJ, Cohen SE, Walker GC. 2008. The SOS Regulatory Network. EcoSal Plus 3. doi:10.1128/ecosalplus.5.4.3.

13. Mandal TN, Mahdi AA, Sharples GJ, Lloyd RG. 1993. Resolution of Holliday intermediates in recombination and DNA repair: Indirect suppression of *ruvA, ruvB*, and *ruvC* mutations. J Bacteriol 175:4325–4334.

14. Sharples GJ, Chan SN, Mahdi AA, Whitby MC, Lloyd RG. 1994. Processing of intermediates in recombination and DNA repair: Identification of a new endonuclease that specifically cleaves Holliday junctions. EMBO J 13:6133–6142.

15. Michel B, Sinha AK, Leach DRF. 2018. Replication Fork Breakage and Restart in *Escherichia coli*. Microbiol Mol Biol Rev 82:e00013–18.

16. Gabbai CB, Marians KJ. 2010. Recruitment to Stalled Replication Forks of the PriA DNA Helicase and Replisome-loading Activities is Essential for Survival. DNA Repair 9:202–209.

17. Michel B, Sandler SJ. 2017. Replication restart in bacteria. J Bacteriol 199:1–13.

18. Heller RC, Marians KJ. 2006. Replisome assembly and the direct restart of stalled replication forks. Nat Rev Mol Cell Biol. 7:932–943.

19. Huang YH, Huang CY. 2014. Structural insight into the DNA-binding mode of the primosomal proteins PriA, PriB, and AnaT. Biomed Res Int http://dx.doi.org/10.1155/2014/195162.

20. Sandler SJ, Leroux M, Windgassen TA, Keck JL. 2021. *Escherichia coli* K- 12 has two distinguishable PriA-PriB replication restart pathways. Mol Microbiol 116:1140–1150.

21. Cox MM. 2007. Regulation of bacterial RecA protein function. Crit Rev Biochem Mol Biol. Crit Rev Biochem Mol Biol 42:41–63.

22. Veaute X, Delmas S, Selva M, Jeusset J, Le Cam E, Matic I, Fabre F, Petit MA. 2005. UvrD helicase, unlike Rep helicase, dismantles RecA nucleoprotein filaments in *Escherichia coli*. EMBO J 24:180–189.

23. Centore RC, Sandler SJ. 2007. UvrD limits the number and intensities of RecA-green fluorescent protein structures in *Escherichia coli* K-12. J Bacteriol 189:2915–2920.

24. Magner DB, Blankschien MD, Lee JA, Pennington JM, Lupski JRR, Rosenberg SM. 2007. RecQ Promotes Toxic Recombination in Cells Lacking Recombination Intermediate-Removal Proteins. Mol Cell 26:273–286.

25. Zhang J, Mahdi AA, Briggs GS, Lloyd RG. 2010. Promoting and avoiding recombination: Contrasting activities of the *Escherichia coli* RuvABC Holliday junction resolvase and RecG DNA translocase. Genetics 185:23–37.

26. Florés MJ, Sanchez N, Michel B. 2005. A fork-clearing role for UvrD. Mol Microbiol 57:1664–1675.

27. Leela JK, Syeda AH, Anupama K, Gowrishankar J. 2013. Rho-dependent transcription termination is essential to prevent excessive genome-wide R- loops in *Escherichia coli*. Proc Natl Acad Sci U S A 110:258–263.

28. Raghunathan N, Kapshikar RM, Leela JK, Mallikarjun J, Bouloc P, Gowrishankar J. 2018. Genome-wide relationship between R-loop formation and antisense transcription in *Escherichia coli*. Nucleic Acids Res 46:3400–3411.

29. Grylak-Mielnicka A, Bidnenko V, Bardowski J, Bidnenko E. Transcription termination factor Rho: a hub linking diverse physiological processes in bacteria. 10.1099/mic.0.000244.

30. Gowrishankar J, Harinarayanan R. 2004. Why is transcription coupled to translation in bacteria? Mol Microbiol 54:598–603.

31. Peters JM, Mooney RA, Grass JA, Jessen ED, Tran F, Landick R. 2012. Rho and NusG suppress pervasive antisense transcription in *Escherichia coli*. Genes Dev 26:2621–2633.

32. Washburn RS, Gottesman ME. 2011. Transcription termination maintains chromosome integrity. Proc Natl Acad Sci U S A 108:792–797.

33. Harinarayanan R, Gowrishankar J. 2003. Host factor titration by chromosomal R-loops as a mechanism for runaway plasmid replication in transcription termination-defective mutants of *Escherichia coli*. J Mol Biol 332:31–46.

34. Gowrishankar J, Krishna Leela J, Anupama K. 2013. R-loops in bacterial transcription: Their causes and consequences. Transcription 4:153–157.

35. Mechulam Y, Blanquet S, Schmitt E. 2013. Translation Initiation. EcoSal Plus 4:1–28.

36. North SH, Kirtland SE, Nakai H. 2007. Translation factor IF2 at the interface of transposition and replication by the PriA-PriC pathway. Mol Microbiol 66:1566–1578.

37. Madison KE, Abdelmeguid MR, Jones-Foster EN, Nakai H. 2012. A new role for translation initiation factor 2 in maintaining genome integrity. PLoS Genet 8(4): e1002648.

38. Madison KE, Jones-Foster EN, Vogt A, Kirtland Turner S, North SH, Nakai H. 2014. Stringent response processes suppress DNA damage sensitivity caused by deficiency in full-length translation initiation factor 2 or PriA helicase. Mol Microbiol 92:28–46.

39. Lee JH, Choi SK, Roll-Mecak A, Burley SK, Dever TE. 1999. Universal conservation in translation initiation revealed by human and archaeal homologs of bacterial translation initiation factor IF2. Proc Natl Acad Sci U S A 96:4342–4347.

40. Howe JG, Hershey JWB. 1984. The rate of evolutionary divergence of initiation factors IF2 and IF3 in various bacterial species determined quantitatively by immunoblotting 140:187–192.

41. Gaur R, Grasso D, Datta PP, Krishna PDV, Das G, Spencer A, Agrawal RK, Spremulli L, Varshney U. 2008. A Single Mammalian Mitochondrial Translation Initiation Factor Functionally Replaces Two Bacterial Factors. Mol Cell 29:180–190.

42. Plumbridge JA, Deville F, Sacerdot C, Petersen HU, Cenatiempo Y, Cozzone A, Grunberg-Manago M, Hershey JW. 1985. Two translational initiation sites in the *infB* gene are used to express initiation factor IF2 alpha and IF2 beta in *Escherichia coli*. EMBO J 4:223–229.

43. Nyengaard NR, Mortensen KK, Lassen SF, Hershey JWB, Sperling-Petersen HU. 1991. Tandem translation of *E.coli* initiation factor IF2β: Purification and characterization in vitro of two active forms. Biochem Biophys Res Commun 181:1572–1579.

44. Laalami S, Putzer H, Plumbridge JA, Grunberg-Manago M. 1991. A severely truncated form of translational initiation factor 2 supports growth of *Escherichia coli*. J Mol Biol 220:335–349.

45. Sacerdot C, Vachon G, Laalami S, Morel-Deville F, Cenatiempo Y, Grunberg-Manago M. 1992. Both forms of translational initiation factor IF2 (α and β) are required for maximal growth of *Escherichia coli.* Evidence for two translational initiation codons for IF2β. J Mol Biol 225:67–80.

46. Mallikarjun J, Gowrishankar J. 2021. Essential role for an isoform of *Escherichia coli* translation initiation factor IF2 in repair of two-ended DNA double-strand breaks. Accompanying article.

47. Anupama K, Leela JK, Gowrishankar J. 2011. Two pathways for RNase E action in *Escherichia coli* in vivo and bypass of its essentiality in mutants defective for Rho-dependent transcription termination. Mol Microbiol 82:1330–1348.

48. Raghunathan N, Goswami S, Leela JK, Pandiyan A, Gowrishankar J. 2019. A new role for *Escherichia coli* Dam DNA methylase in prevention of aberrant chromosomal replication. Nucleic Acids Res 47:5698–5711.

49. Ali N, Gowrishankar J. 2020. Cross-subunit catalysis and a new phenomenon of recessive resurrection in *Escherichia coli* RNase E. Nucleic Acids Res 48:847–861.

50. Leela JK, Raghunathan N, Gowrishankar J. 2021. Topoisomerase I Essentiality, DnaA-Independent Chromosomal Replication, and Transcription-Replication Conflict in *Escherichia coli*. J Bacteriol 203: e00195–21

51. Johnson DBF, Wang C, Xu J, Schultz MD, Schmitz RJ, Ecker JR, Wang L. 2012. Release factor one is nonessential in *Escherichia coli*. ACS Chem Biol 7:1337–1344.

52. Hu K, Artsimovitch I. 2017. A screen for *rfaH* suppressors reveals a key role for a connector region of termination factor *rho*. mBio 8(3): e00753–17.

53. McGlynn P, Lloyd RG. 2000. Modulation of RNA polymerase by (p)ppGpp reveals a RecG-dependent mechanism for replication fork progression. Cell 101:35–45.

54. Trautinger BW, Jaktaji RP. 2005. RNA Polymerase Modulators and DNA Repair Activities Resolve Conflicts between DNA Replication and Transcription. Mol Cell 19:247–258.

55. Kamarthapu V, Epshtein V, Benjamin B, Proshkin S, Mironov A, Cashel M, Nudler E. 2016. ppGpp couples transcription to DNA repair in *E. coli*. Science 352:993–996.

56. Carles-Kinch K, George JW, Kreuzer KN. 1997. Bacteriophage T4 UvsW protein is a helicase involved in recombination, repair and the regulation of DNA replication origins. EMBO J 16:4142–4151.

57. Dudas KC, Kreuzer KN. 2001. UvsW Protein Regulates Bacteriophage T4 Origin-Dependent Replication by Unwinding R-Loops. Mol Cell Biol 21:2706–2715.

58. Brandi A, Piersimoni L, Feto NA, Spurio R, Alix JH, Schmidt F, Gualerzi CO. 2019. Translation initiation factor IF2 contributes to ribosome assembly and maturation during cold adaptation. Nucleic Acids Res 47:4652–4662.

59. Xia J, Chen LT, Mei Q, Ma CH, Halliday JA, Lin HY, Magnan D, Pribis JP, Fitzgerald DM, Hamilton HM, Richters M, Nehring RB, Shen X, Li L, Bates D, Hastings PJ, Herman C, Jayaram M, Rosenberg SM. 2016. Holliday junction trap shows how cells use recombination and a junction-guardian role of RecQ helicase. Sci Adv 2: e1601605.

60. Boguslawski SJ, Smith DE, Michalak MA, Mickelson KE, Yehle CO, Patterson WL, Carrico RJ. 1986. Characterization of monoclonal antibody to DNA · RNA and its application to immunodetection of hybrids. J Immunol Methods 89:123–130.

61. Lovett ST, Hurley RL, Sutera VA, Aubuchon RH, Lebedeva MA. 2002. Crossing over between regions of limited homology in *Escherichia coli*: RecA-dependent and RecA-independent pathways. Genetics 160:851–859.

62. Dutra BE, Sutera VA, Lovett ST. 2007. RecA-independent recombination is efficient but limited by exonucleases. Proc Natl Acad Sci U S A 104:216–221.

63. Laban A, Cohen A. 1981. Interplasmidic and intraplasmidic recombination in *Escherichia coli* K-12. MGG Mol Gen Genet 184:200–207.

64. Cohen A, Laban A. 1983. Plasmidic recombination in *Escherichia coli* K-12: The role of *recF* gene function. MGG Mol Gen Genet 189:471–474.

65. Konrad EB. 1977. Method for the isolation of *Escherichia coli* mutants with enhanced recombination between chromosomal duplications. J Bacteriol 130:167–172.

66. Zhang G, Deng E, Baugh L, Kushner SR. 1998. Identification and Characterization of *Escherichia coli* DNA Helicase II Mutants That Exhibit Increased Unwinding Efficiency. J Bacteriol 180:377–387.

67. Nazir A, Harinarayanan R. 2016. Inactivation of cell division protein FtsZ by SulA makes Lon indispensable for the viability of a ppGpp^0^ strain of *Escherichia coli*. J Bacteriol 198:688–700.

68. Brüning JG, Marians KJ. 2021. Bypass of complex co-directional replication-transcription collisions by replisome skipping. Nucleic Acids Res 49:9870–9885.

69. Seigneur M, Ehrlich SD, Michel B. 2000. RuvABC-dependent double-strand breaks in dnaB^ts^ mutants require RecA. Mol Microbiol 38:565–574.

70. Mahdi AA, Buckman C, Harris L, Lloyd RG. 2006. Rep and PriA helicase activities prevent RecA from provoking unnecessary recombination during replication fork repair. Genes Dev 20:2135–2147.

71. Khan SR, Kuzminov A. 2012. Replication forks stalled at ultraviolet lesions are rescued via RecA and RuvABC protein-catalyzed disintegration in *Escherichia coli*. J Biol Chem 287:6250–6265.

72. Krejci L, Altmannova V, Spirek M, Zhao X. 2012. Homologous recombination and its regulation. Nucleic Acids Res 40:5795–5818.

73. Matic I, Rayssiguier C, Radman M. 1995. Interspecies Gene Exchange in Bacteria: The Role of SOS and Mismatch Repair Systems in Evolution of Species. Cell 80:507–515.

74. Renzette N, Gumlaw N, Sandler SJ. 2007. DinI and RecX modulate RecA-DNA structures in *Escherichia coli* K-12. Mol Microbiol 63(1):103–115.

75. Miller JH. 1992. A Short Course in Bacterial Genetics: A Laboratory Manual and Handbook for *Escherichia coli* and Related Bacteria. Cold Spring Harb Lab Press NY.

76. Baba T, Ara T, Hasegawa M, Takai Y, Okumura Y, Baba M, Datsenko KA, Tomita M, Wanner BL, Mori H. 2006. Construction of *Escherichia coli* K-12 in-frame, single-gene knockout mutants: The Keio collection. Mol Syst Biol 2:2006.0008.

77. Bolivar F, Rodriguez RL, Greene PJ, Betlach MC, Heyneker HL, Boyer HW, Crosa JH, Falkow S. 1977. Construction and characterization of new cloning vehicle. II. A multipurpose cloning system. Gene 2:95–113.

78. Chang ACY, Cohen SN. 1978. Construction and characterization of amplifiable multicopy DNA cloning vehicles derived from the p15A cryptic miniplasmid. J Bacteriol 134:1141–1156.

79. Lerner CG, Inouye M. 1990. Low copy number plasmids for regulated low-level expression of cloned genes in *Escherichia coli* with blue-white insert screening capability. Nucleic Acids Res. Nucleic Acids Res 18:4631.

80. Andrews AE, Lawley B, Pittard AJ. 1991. Mutational analysis of repression and activation of the *tyrP* gene in *Escherichia coli*. J Bacteriol 173:5068–5078.

81. Amann E, Ochs B, Abel KJ. 1988. Tightly regulated *tac* promoter vectors useful for the expression of unfused and fused proteins in *Escherichia coli*. Gene 69:301–315.

82. Datsenko KA, Wanner BL. 2000. One-step inactivation of chromosomal genes in *Escherichia coli* K-12 using PCR products. Proc Natl Acad Sci U S A 97:6640–6645.

83. Gowrishankar J. 1985. Identification of osmoresponsive genes in *Escherichia coli*: Evidence for participation of potassium and proline transport systems in osmoregulation. J Bacteriol 164:434–445.

84. Sambrook D, Russell J. 2001. Molecular Cloning: A Laboratory Manual. 3rd edn. Cold Spring Harb Lab Press NY.

85. Boyd D, Weiss DS, Chen JC, Beckwith J. 2000. Towards single-copy gene expression systems making gene cloning physiologically relevant: Lambda InCh, a simple *Escherichia coli* plasmid-chromosome shuttle system. J Bacteriol 182:842–847.

