## Supplementary Data including Supplementary Text, Supplementary References, Supplementary Table S1, and Supplementary Figures S1-S5. for "Modulation of RecFORQ- and RecA-mediated homologous recombination in *Escherichia coli* by isoforms of translation initiation factor IF2"

**CONTENTS:**

**Supplementary Text.**

**Supplementary References.**

**Supp. Table S1.** List of *E. coli* strains.

**Supp. Fig. S1.** Features associated with *rho*-136$^{opal}$ mutation.

**Supp. Fig. S2.** Blue-white assays at 30°, with *rho*$^{+}$ *infB*$^{+}$ shelter plasmid pHYD5212, for *rho*-136$^{opal}$ *ruv* synthetic lethality and its suppression.

**Supp. Fig. S3.** IF2 isoforms and *infB*-161$^{ochre}$ mutation.

**Supp. Fig. S4.** Modulation of *rho ruv* synthetic lethality by differential expression of IF2 isoforms.

**Supp. Fig. S5.** UV tolerance and SOS induction in strains expressing different IF2 isoforms.

### Supplementary Text

**Details of plasmids constructed.** ***(a) pHYD5207 and pHYD5208*** (pTrc99A derivatives expressing, respectively, IF2-2,3 and IF2-3): Plasmids pHYD5207 and pHYD5208 were each constructed as an intermediate step in obtaining ectopic chromosomal integration of $P_{trc} - \Delta\mathit{1}$ (expressing IF2-2,3) and $P_{trc} - \Delta\mathit{1,2}$ (expressing IF2-3), respectively, by the method of Boyd et al. (1). Primer pair 5’-ACGC*GTCGAC*GCGAAACGTGAAGCTGCGGAA-3’ and 5’-TCC*AAGCTT*CATGGAACCCTAAAAACCTTA-3’ was used for PCR with genomic DNA from strains GJ13519 (for pHYD5207) or GJ15445 (for pHYD5208) as template (*SalI* and *HindIII* sites, respectively, in the two primers italicized). Each PCR amplicon was digested with *SalI* and *HindIII* and cloned into the *SalI-HindIII* sites of pTrc99A, to generate pHYD5207 and pHYD5208, respectively. The primers are designed to amplify a part of the *infB* ORF from codons 148 to 890 (that is, to the end of the gene), so that the initiation codons for both IF2-2 and IF2-3 are contained in the PCR amplicon; however, since strain GJ15445 carries the *ochre* mutation in codon 161 of *infB*, the amplicon therefrom is disabled from expressing IF2-2.

***(b) pHYD5212*** (pMU575-derived shelter plasmid carrying *rho*$^+$ and *infB*$^+$): Plasmid pHYD5212 is a derivative of the vector pMU575 and was constructed in three steps as follows. In the first step, a 7.9-kb *HindIII-SalI* fragment from the *E. coli* genome carrying the genes *rbfA',* *infB, nusA, yhbC, metY, argG,* and *'yhbX* was subcloned, through an intermediate vector, into the corresponding sites of pMU575, to generate plasmid pHYD2906. In the second step, the *rho*$^+$ gene was PCR-amplified from genomic DNA template of *Salmonella enterica* LT2 with the primer pair 5’-TATCCATACAAAGACATATTGTCGACCGCAAACAATGTGTATTGTG-3’ and 5’-AATTAATTCCCGATCTGCAGGTCGACTACGTTATTCTTAAATTGTCAGG-3’. The PCR amplicon was cloned into *SalI*-digested pHYD2906 by the Gibson assembly protocol (2), so as to generate plasmid pHYD5211; the *rho*$^+$ gene on pHYD5211 was sequence verified. Finally, the *argG* gene on pHYD5211 was disrupted with the aid of the pKD46-based recombineering method of Datsenko and Wanner (3), by deletion and insertion of the Sp$^R$ gene sourced from plasmid pCL1920, so as to generate plasmid pHYD5212; for this purpose, pCL1920 DNA was used as template for PCR with primer pair 5’-TCTCCCGGTAGGTCAACGTAGTTCGTAAGCTGTAATGCAAG-3’ and 5’-GATTCTCCACCTGCGGCACGCCTGATAGTTTGGCTGTGAG-3’, and the amplicon so obtained was subjected to a second PCR with primer pair 5'-ATGACGACGATTCTCAAGCATCTCCCGGTAGGTCAACGTA-3' and 5'-TTACTGGCCTTTGTTTTCCAGATTCTCCACCTGCGGCACG-3' to obtain the DNA used for recombineering.

***(c) pHYD5215 and pHYD5216*** (derivatives of plasmids pBR322 and pACYC184, respectively, with non-overlapping deletions within *tetA* gene): Plasmids pHYD5215 and pHYD5216 were constructed for use in the inter-plasmid recombination assay. The former is a pBR322 derivative with deletion of the *tetA* promoter region between *EcoRI* and *HindIII* sites, and was generated by digestion of pBR322 with the two enzymes, end-filling by Klenow fragment of DNA polymerase I, and recircularization by ligation. The latter, which is a pACYC184 derivative with deletion of the *tetA* gene between *BamHI* and *SalI* sites, was generated by a similar approach mutatis mutandis.

**HR assays.** ***(a) Konrad assay***: Strain SK707 or its derivatives were each grown from a small inoculum (around $10^4$ cells) to stationary phase in LB. The cells were washed once in minimal A buffer, resuspended in minimal A, and plated at appropriate dilutions (i) on LB to determine viable counts, and (ii) on 0.2% lactose-minimal A plates that had been pre-spread with $10^9$ cells of the Δ*lac* strain GJ15410 as scavenger (4), to determine the number $Lac^+$ recombinants.

***(b) Inter-plasmid recombination assay***: Strains transformed with both plasmids pHYD5215 and pHYD5216 were each grown from a small inoculum (around $10^4$ cells) to stationary phase in LB supplemented with Amp and Cm. Suitable dilutions were plated on LB-Amp Cm and LB-Amp Cm Tet to determine viable counts and the number of $Tet^R$ recombinants, respectively.

***(c) Conjugation***: The Hfr donor in conjugation GJ19848 was $Tet^R$ *metB leuD* $Cm^S$ while the recipients GJ19849 or GJ19850 were both $Tet^S$ *leuA* $Cm^R$. Donor and recipient cultures, grown to early exponential phase in LB, were mixed at 1:10 ratio and incubated for 1 hour. The cells were washed twice in minimal A buffer, and $Tet^R$ exconjugants were selected on glucose-minimal A supplemented with Tet, Cm and the amino acids Met, Ile, Leu and Val. $Tet^R$ $Leu^+$ exconjugants were selected on the same medium but with the branched-chain amino acid supplements omitted. Exconjugant frequencies were calculated per viable Hfr donor.

***(d) Phage P1 transductions***: Phage P1 transduction frequencies were determined as the mean of around twenty independent experiments for each genotype encoding a different IF2 isoform with selections that included the following: (i) donor marker selected in transduction: Δ*yihF*::Kan, Δ*serA*::Kan, Δ*ybfP*::Kan, Δ*ilvA*::Kan, Δ*leuA*::Kan, Δ*leuD*::Kan, Δ*thrA*::Kan, and Δ*racC*::Kan; and (ii) recipient marker replaced in selection to prototrophy: Δ*serA*::Kan and Δ*ilvA*::Kan. Transduction frequency in each experiment was calculated per plaque-forming unit of phage.

**Replacement of Cm$^R$ cassette with Kan$^R$ cassette by recombineering.** To replace the Cm$^R$ gene in each of the Nakai ectopic *infB* constructs with Kan$^R$, the Datsenko and Wanner recombineering method (3) with plasmid pKD46 was used. The Kan$^R$ cassette was amplified (with flanking FRT sites) from plasmid pKD13 using the primer pair: 5'-TCACTGGATATACCACCGTTGATATATCCCAATGGCATCGATTCCGGGGATCCGTCGACC-3' and 5'-GCATTCTGCCGACATGGAAGCCATCACAGACGGCATGATGTGTAGGCTGGAGCTGCTTCG-3'.

### Supplementary References


1. Boyd D, Weiss DS, Chen JC, Beckwith J. 2000. Towards single-copy gene expression systems making gene cloning physiologically relevant: Lambda InCh, a simple *Escherichia coli* plasmid- chromosome shuttle system. J Bacteriol 182:842–847.

2. Gibson DG, Young L, Chuang RY, Venter JC, Hutchison CA, Smith HO. Enzymatic assembly of DNA molecules up to several hundred kilobases. Nat Meth 6(5):343-347.

3. Datsenko KA, Wanner BL. 2000. One-step inactivation of chromosomal genes in *Escherichia coli* K-12 using PCR products. Proc Natl Acad Sci U S A 97:6640–6645.

4. Reddy M, Gowrishankar J. 2000. Characterization of the *uup* locus and its role in transposon excisions and tandem repeat deletions in *Escherichia coli*. J Bacteriol 182:1978–1986.

5. Singer M, Baker TA, Schnitzler G, Deischel SM, Goel M, Dove W, Jaacks KJ, Grossman AD, Erickson JW, Gross CA. 1989. A collection of strains containing genetically linked alternating antibiotic resistance elements for genetic mapping of *Escherichia coli*. Microbiol Rev 53:1–24.

6. Zhang G, Deng E, Baugh L, Kushner SR. 1998. Identification and Characterization of *Escherichia coli* DNA Helicase II Mutants That Exhibit Increased Unwinding Efficiency. J Bacteriol 180:377-387.

7. Raghunathan N, Kapshikar RM, Leela JK, Mallikarjun J, Bouloc P, Gowrishankar J. 2018.

Genome-wide relationship between R-loop formation and antisense transcription in *Escherichia coli*. Nucleic Acids Res 46:3400–3411.

8. Leela JK, Syeda AH, Anupama K, Gowrishankar J. 2013. Rho-dependent transcription termination is essential to prevent excessive genome-wide R-loops in *Escherichia coli*. Proc Natl Acad Sci U S A 110:258–263.

9. Sanyal R, Singh V, Harinarayanan R. 2019. A novel gene contributing to the initiation of fatty acid biosynthesis in *Escherichia coli*. J Bacteriol 201:1–20.

10. Nazir A, Harinarayanan R. 2016. Inactivation of cell division protein FtsZ by SulA makes Lon indispensable for the viability of a ppGpp$^0$ strain of *Escherichia coli*. J Bacteriol 198:688–700.

11. Harinarayanan R, Gowrishankar J. 2003. Host factor titration by chromosomal R-loops as a mechanism for runaway plasmid replication in transcription termination-defective mutants of *Escherichia coli*. J Mol Biol 332:31–46.

12. Layton JC, Foster PL. 2003. Error-prone DNA polymerase IV is controlled by the stress-response sigma factor, RpoS, in *Escherichia coli*. Mol Microbiol 50:549–561.

13. Seigneur M, Bidnenko V, Ehrlich SD, Michel B. 1998. RuvAB acts at arrested replication forks. Cell 95:419–430.

14. Gaur R, Grasso D, Datta PP, Krishna PDV, Das G, Spencer A, Agrawal RK, Spremulli L, Varshney U. 2008. A Single Mammalian Mitochondrial Translation Initiation Factor Functionally Replaces Two Bacterial Factors. Mol Cell 29:180–190.

15. Raghunathan N, Goswami S, Leela JK, Pandiyan A, Gowrishankar J. 2019. A new role for *Escherichia coli* Dam DNA methylase in prevention of aberrant chromosomal replication. Nucleic Acids Res 47:5698–5711.

16. Carles-Kinch K, George JW, Kreuzer KN. 1997. Bacteriophage T4 UvsW protein is a helicase involved in recombination, repair and the regulation of DNA replication origins. EMBO J 16:4142–4151.

17. Anupama K, Leela JK, Gowrishankar J. 2011. Two pathways for RNase E action in *Escherichia coli* in vivo and bypass of its essentiality in mutants defective for Rho-dependent transcription termination. Mol Microbiol 82:1330–1348.

18. Baba T, Ara T, Hasegawa M, Takai Y, Okumura Y, Baba M, Datsenko KA, Tomita M, Wanner BL, Mori H. 2006. Construction of *Escherichia coli* K-12 in-frame, single-gene knockout mutants: The Keio collection. Mol Syst Biol 2:2006.0008.

19. Saisree L, Reddy M, Gowrishankar J. 2000. lon incompatibility associated with mutations causing SOS induction: Null uvrD alleles induce an SOS response in *Escherichia coli*. J Bacteriol 182:3151–3157.

20. Xia J, Chen LT, Mei Q, Ma CH, Halliday JA, Lin HY, Magnan D, Pribis JP, Fitzgerald DM, Hamilton HM, Richters M, Nehring RB, Shen X, Li L, Bates D, Hastings PJ, Herman C,

Jayaram M, Rosenberg SM. 2016. Holliday junction trap shows how cells use recombination and a junction-guardian role of RecQ helicase. Sci Adv 2: e1601605.

21. Mandal TN, Mahdi AA, Sharples GJ, Lloyd RG. 1993. Resolution of Holliday intermediates in recombination and DNA repair: Indirect suppression of *ruvA*, *ruvB*, and *ruvC* mutations. J Bacteriol 175:4325–4334.

22. Madison KE, Abdelmeguid MR, Jones-Foster EN, Nakai H. 2012. A new role for translation initiation factor 2 in maintaining genome integrity. PLoS Genet 8(4): e1002648.

23. Saroja GN, Gowrishankar J. 1996. Roles of SpoT and FNR in $NH4^{+}$ assimilation and osmoregulation in GOGAT (glutamate synthase)-deficient mutants of *Escherichia coli*. J Bacteriol 178:4105–4114.

**Table S1.** List of *E. coli* K-12 strains

| **Strain** [a] | **Genotype** [b] |
|---|---|
| **MG1655** | *E. coli* K-12 wild-type |
| **CAG5052** | Hfr(PO3) *relA1 spoT1 metB1 btuB3191*::Tn*10* |
| **SK707** | *lacMS286* Φ80dII*lac*BK1 *argH1 hisG4 ilvD188 metE46 rpsL* |
| **GJ13519** | MG1655 Δ(*argF-lac)U169* (Φ80 lysogen) |
| **GJ13567**[c] engineered | MG1655 Δ(*argF-lac)U169* Δ*rho*::Kan *rpoB**35 *btuB*::Tn*10*, along with 42 deletions in genome |
| **GJ14818** | GJ13519 *rho-4* |
| **GJ15396**[c] | GJ13519 *rho*-136$^{opal}$ |
| **GJ15410** | MG1655 Δ*lacIZYA*::FRT *galEp3 sulA11* |
| **GJ15429**[c] | GJ13519 *rho*-136$^{opal}$ *prfB*$^{+}$ (from *E. coli* B) |
| **GJ15441**[d] | GJ15410 *rho*-136$^{opal}$ |
| **GJ15445**[d] | GJ15410 *rho*-136$^{opal}$ *infB*-161$^{ochre}$ *zhc-914*::Tn*10*dKan |
| **GJ15446**[d] | GJ15410 *rho*-136$^{opal}$ Δ*ruvABC::*Cm *infB*-161$^{ochre}$ |
| **GJ15447**[d] | GJ15410 *rho*-136$^{opal}$ Δ*ruvABC::*Cm |
| **GJ15449**[d] | GJ15410 *infB*-161$^{ochre}$ |
| **GJ15457**[d] | GJ15447 *att* λ::<*lacI* ($P_{trc}$ – *infB*Δ*1,2*)-Amp> |
| **GJ15458**[d] | GJ15457 Δ*infB*::Kan |
| **GJ15460**[d] | GJ15447 Δ*rac ydaH*::Tn*10* Δ*recA*::Kan |

| | |
|---|---|
| **GJ15464**[d] | GJ15447 Δ*infB*::Kan *att* λ::<*lacI* ($P_{trc}$ – *infB*Δ*1*)-Amp> |
| **GJ15465**[d] | GJ15447 Δ*rac ydaH*::Tn*10 att* λ::<*lacI* ($P_{tac}$ – UvsW)-Amp> |
| **GJ15466**[d] | GJ15447 Δ*rac ydaH*::Tn*10 att* λ::<*lacI* ($P_{tac}$ – UvsW-K141R)-Amp> |
| **GJ15471**[d] | GJ15447 Δ*rac ydaH*::Tn*10* Δ*recO*::Kan |
| **GJ15475**[d] | GJ15410 Δ*infB*::Kan *att* λ::<*lacI* ($P_{trc}$ – *infB*Δ*1*)-Amp> |
| **GJ15485**[d] | GJ15441 Δ*rac ydaH*::Tn*10* Δ*ruvA*::FRT *lexA3 malB*::Tn*9* |
| **GJ15487**[d] | GJ15447 Δ*rac ydaH*::Tn*10* Δ*recB*::Kan |
| **GJ15488** | GJ15410 Δ*ruvA*::FRT |
| **GJ15490**[d] | GJ15441 Δ*rac ydaH*::Tn*10* Δ*ruvA*::FRT Δ*recA*::Kan |
| **GJ15491**[d] | GJ15447 Δ*rac ydaH*::Tn*10* Δ*recR*::Kan |
| **GJ15494** | GJ15410 Δ*infB*::FRT *flgJ*::<*nusA infB*(Δ*1*)-Cm> |
| **GJ15497**[d] | GJ15447 Δ*rac ydaH*::Tn*10* Δ*recQ*::Kan |
| **GJ15498**[d] | GJ15441 Δ*rac ydaH*::Tn*10* Δ*ruvA*::FRT Δ*infB*::Kan *flgJ*::<*nusA infB*(*wt*)-Cm> |
| **GJ15499**[d] | GJ15441 Δ*rac ydaH*::Tn*10* Δ*ruvA*::FRT Δ*infB*::Kan *flgJ*::<*nusA infB*(Δ*1*)-Cm> |
| **GJ15500**[d] | GJ15441 Δ*rac ydaH*::Tn*10* Δ*ruvA*::FRT Δ*infB*::Kan *flgJ*::<*nusA infB*(Δ*2,3*)-Cm> |
| **GJ19101** | GJ15410 Δ*infB*::Kan *flgJ*::<*nusA infB*(*wt*)-Cm> |
| **GJ19102** | GJ15410 Δ*infB*::Kan *flgJ*::<*nusA infB(*Δ*2,3*)-Cm> |
| **GJ19125**[d] | GJ15441 Δ*infB*::Kan *att*λ::<*lacI* ($P_{trc}$ – *infB*Δ*1*)-Amp> |
| **GJ19127** | GJ15410 Δ*attλ*::<($P_{N25}$-*tetR*)-FRT> *zfe-2512.7*::<($P_{N25tetO}$-RDG)-FRT> |
| **GJ19131**[d] | GJ15410Δ*infB*::FRT *rho-4* Δ*ruvC*::Kan *flgJ*::<*nusA infB(*Δ*2,3*)-Cm> |
| **GJ19132**[d] | GJ19131 *att* λ::<*lacI* ($P_{trc}$ – *infB*Δ*1*)-Amp> |
| **GJ19134**[d] | GJ15410 Δ*infB*::FRT *rho*-4 Δ*ruvC*::Kan *flgJ*::<*nusA infB*(*wt*)-Cm> |
| **GJ19150**[d] | GJ15441 Δ*ruvC*::FRT |
| **GJ19152**[d] | GJ19150 Δ*infB*::Kan *att* λ::<*lacI* ($P_{trc}$ – *infB*Δ*1*)-Amp> |
| **GJ19161** | GJ19127 Δ*uvrD*::FRT |
| **GJ19162** | SK707 Δ*uvrD*::FRT |
| **GJ19165** | GJ19162 Δ*infB*::Kan *flgJ*::<*nusA infB*(Δ*1*)-Cm> |
| **GJ19166** | GJ15494 Δ*ruvA*::FRT |
| **GJ19171** | SK707 Δ*infB*::Kan *flgJ*::<*nusA infB(*Δ*2,3*)-Cm> |

**GJ19181** SK707 *rho*-136$^{opal}$

**GJ19184** GJ19162 Δ*infB*::Kan *flgJ*::<*nusA infB(*Δ*2,3*)-Cm>

**GJ19186** SK707 Δ*infB*::Kan *flgJ*::<*nusA infB*(Δ*1*)-Cm>

**GJ19193** GJ15410 Δ*infB*::FRT *flgJ*::<*nusA infB*(*wt*)-Cm>

**GJ19194** GJ15410 Δ*infB*::FRT *flgJ*::<*nusA infB(*Δ*2,3*)-Cm>

**GJ19195** GJ15410 Δ*infB*::FRT *flgJ*::<*nusA infB*(Δ*1*)-Kan>

**GJ19196** GJ15410 Δ*infB*::FRT *flgJ*::<*nusA infB(*Δ*2,3*)-Kan>

**GJ19197** GJ15410 Δ*infB*::FRT *flgJ*::<*nusA infB*(*wt*)-Kan>

**GJ19379**[d] GJ15441 *att* λ::<*araC* ($P_{ara}$ – *rho*)-Amp>

**GJ19380**[d] GJ15445 *att* λ::<*araC* ($P_{ara}$ – *rho*)-Amp>

**GJ19381**[d] GJ15447 *att* λ::<*araC* ($P_{ara}$ – *rho*)-Amp>

**GJ19382**[d] GJ15446 *att* λ::<*araC* ($P_{ara}$ – *rho*)-Amp>

**GJ19801** GJ19161 Δ*infB*::Kan *flgJ*::<*nusA infB*(*wt*)-Cm>

**GJ19802** GJ19161 Δ*infB*::Kan *flgJ*::<*nusA infB(*Δ*2,3*)-Cm>

**GJ19803** GJ19161 Δ*infB*::Kan *flgJ*::<*nusA infB*(Δ*1*)-Cm>

**GJ19835** GJ19161 Δ*infB*::FRT Δ*recA*::Kan *flgJ*::<*nusA infB*(*wt*)-Cm>

**GJ19839** MG1655 Δ*lacIZYA*::FRT *galEp*3 Δ*infB*::FRT *flgJ*::<*nusA infB*(*wt*)-Cm> $P_{sulA}$ ::(FRT-*lacZY*-Kan-*oriR6K*-FRT)

**GJ19840** MG1655 Δ*lacIZYA*::FRT *galEp*3 Δ*infB*::FRT *flgJ*::<*nusA infB*(*(*Δ*2,3*)-Cm> $P_{sulA}$ ::(FRT-*lacZY*-Kan-*oriR6K*-FRT)

**GJ19841** MG1655 Δ*lacIZYA*::FRT *galEp*3 Δ*infB*::FRT *flgJ*::<*nusA infB*(Δ*1*)-Cm> $P_{sulA}$ ::(FRT-*lacZY*-Kan-*oriR6K*-FRT)

**GJ19843** GJ19161 Δ*infB*::FRT *flgJ*::<*nusA infB*(*wt*)-Cm>

**GJ19845**[c] GJ15396 Δ*recA*::Kan

**GJ19846**[d] GJ15447 *rpoB*35 btuB*::Tn*10*

**GJ19847** GJ15410 Δ*uvrD*::Kan

**GJ19848** CAG5052 Δ*leuD*::Kan

**GJ19849**[d] GJ19193 *rho*-136$^{opal}$ Δ*leuA*::Kan

**GJ19850** GJ19193 Δ*leuA*::Kan

**GJ19853**[d] GJ15447 *rus-1 purE*::Tn*10*

[a] Strains MG1655 (4), CAG5052 (5), SK707 (6), GJ13519 (7), and GJ13567 (8) have been described earlier; GJ14818 was from our laboratory collection, constructed by J Krishna Leela; all other strains were constructed in the present study.

[b] The following alleles and constructs have been described earlier: Δ*lacIZYA* (9); $P_{sulA}$ ::(FRT-*lacZY*-Kan-*oriR6K*-FRT) (10); *galEp3* and *rho-4* (11); *sulA11* (12); Δ*ruvABC*::Cm (13); Δ*infB*::Kan (14), whose FRT derivative was generated in this study; *rpoB*35 btuB*::Tn*10* (15), *att* λ::<*lacI* ($P_{tac}$ – UvsW)-Amp> and *att* λ::<*lacI* ($P_{tac}$ – UvsW-K141R)-Amp> (16); *att* λ::<*araC* ($P_{ara}$ – *rho*)-Amp> (17); Keio insertion-deletions (::Kan or their corresponding ::FRT derivatives) of *recA*, *recB*, *recO*, *recR*, *recQ*, *ruvA*, *ruvC*, and *uvrD* (18); *lexA3 malB*::Tn*9* (19); *cycA*::Tn*10* (5); Δ*att*λ::<($P_{N25}$-*tetR*)-FRT> and *zfe-2512.7*::<($P_{N25tetO}$-RDG)-FRT> (20); *rus-1* (21); and *flgJ*::<*nusA infB*(*wt*)-Cm>, *flgJ*::<*nus AinfB*((Δ*2,3*)-Cm>, and *flgJ*::<*nusA infB*(Δ*1*)-Cm>, that are designated in the text as Δ*Nil*, Δ*2,3* and Δ*1*, respectively (22). The Δ*rac* allele was sourced from strain AB1157 (23), and it was linked to *ydaH*::Tn*10* (5) (by J Krishna Leela).

[c] The indicated strains were routinely maintained with the *rho*$^{+}$ shelter plasmid pHYD2411.

[d] The indicated strains were routinely maintained with the *rho*$^{+}$ *infB*$^{+}$ shelter plasmid pHYD5212.

**Supp. Fig. S1**

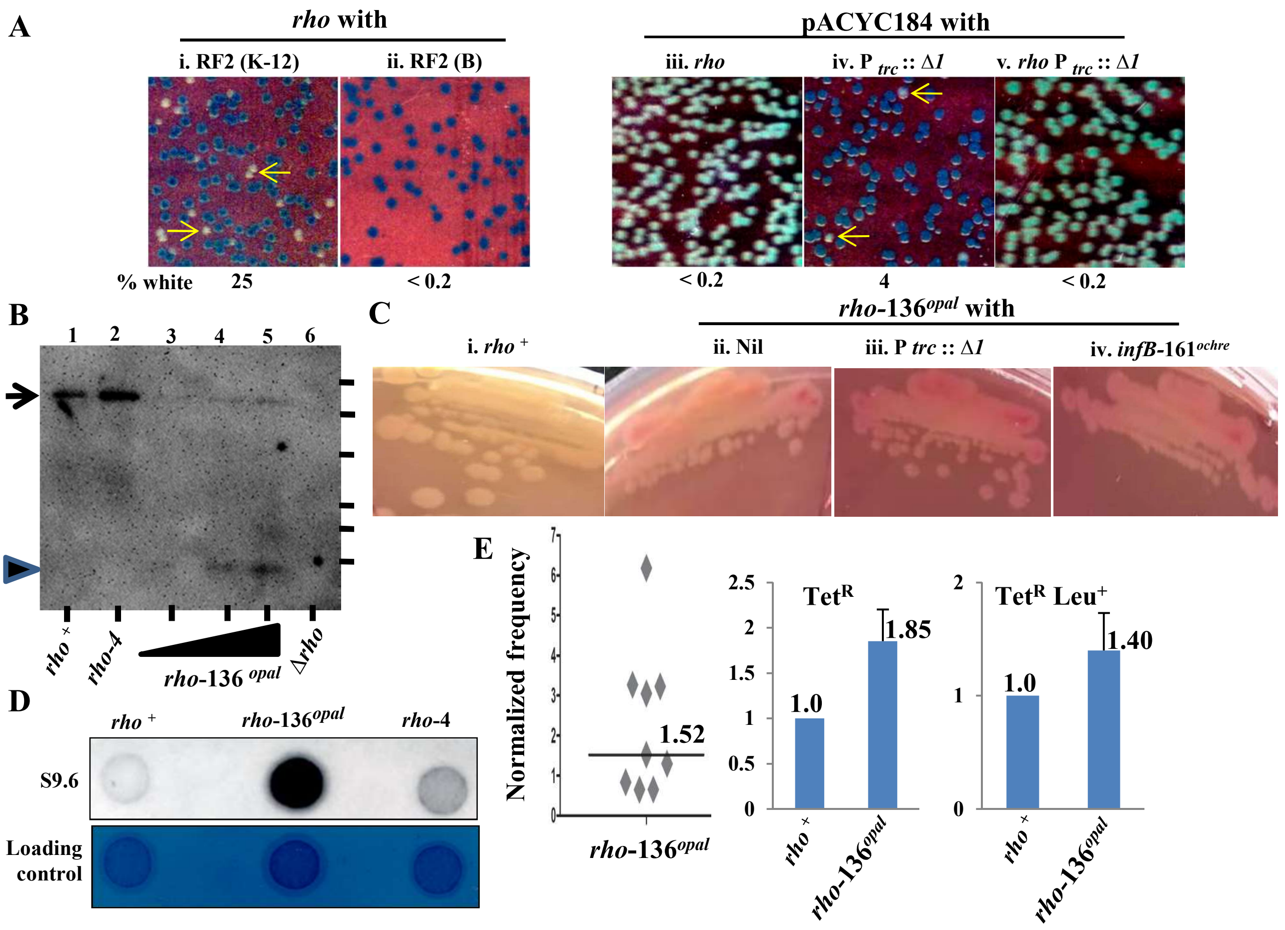


**Supplementary Figure S1**: Features associated with *rho*-136$^{opal}$ mutation. All strain numbers mentioned are prefixed with GJ. **(A)** Blue-white screening assays for strains carrying shelter plasmid pHYD2411 (*rho*$^{+}$, sub-panels i-ii on defined medium) or pHYD5212 (*rho*$^{+}$ *infB*$^{+}$, sub-panels iii-v on LB supplemented with IPTG). Notations are as described in legend to Figure 1A; designation *rho* refers to *rho*-136$^{opal}$, and K-12 and B within parentheses refer to the corresponding *E. coli* sub-species. Strains were derivatives of (with shelter plasmid as indicated, and also with pACYC184 for sub-panels iii-v): i, 15396; ii, 15429; iii, 15441; iv, 15475; and v, 19125. **(B)** Western blotting (following electrophoresis on 10% polyacrylamide gel with sodium dodecyl sulphate) with anti-Rho antibody, for *rho*$^{+}$, *rho-4*, *rho*-136$^{opal}$, and Δ*rho-rpoB*35* strains (13519, 14818, 15396, and 13567, respectively). Bands corresponding to full-length (47 kDa) and truncated (15 kDa) Rho polypeptides are marked by arrow and arrowhead, respectively. Positions of migration of molecular mass standards are depicted by bars at right (from top: 58, 46, 32, 25, 22, and 17 kDa, respectively). **(C)** MacConkey-galactose agar phenotypes for *galEp3* derivatives (plate was also supplemented with IPTG). Strains were (from left): 15410, 15441, 19125, and 15445. **(D)** Immunoblot with S9.6 antibody (top) and methylene blue-stained image of blot (as loading control, bottom) of total nucleic acid preparations from strains *rho*$^{+}$ (15410), *rho*-136$^{opal}$ (15441), and *rho-4* (14818). **(E)** Recombination frequency data from the Konrad assay (left) and conjugation experiments (middle and right), after normalization to the value for the cognate control strains (depicted in Figure 4 for Konrad assay, and within each sub-panel for data from conjugation experiments; the actual values in the latter were $2.5*10^{-3}$ and $5.1*10^{-5}$ per Hfr donor cell for Tet$^{R}$ and Tet$^{R}$ Leu$^{+}$ selections, respectively). Notations for the Konrad assay are as described in legend to Figure 4. Mean values (given alongside each bar) and standard errors are shown for the *rho* mutant in the conjugation experiments. Strains were: for Konrad assay, 19181; and for conjugation experiments, 19848 as Hfr donor and 19850 (*rho*$^{+}$) and 19849 (*rho*-136$^{opal}$) as recipients.

**Supp. Fig. S2**

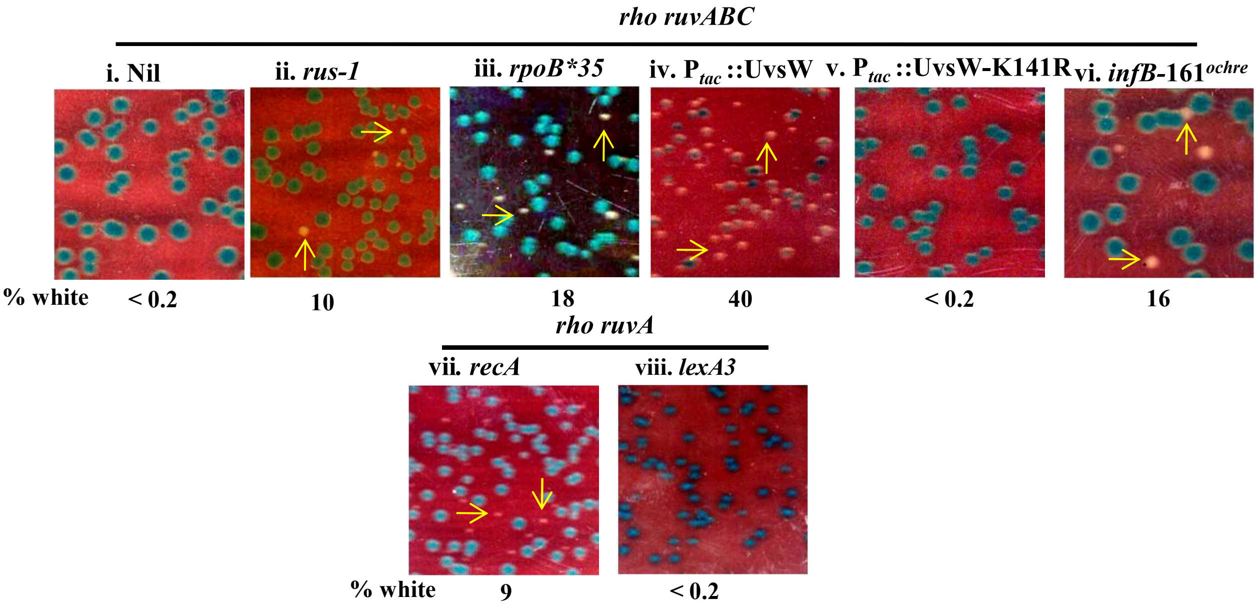


**Supplementary Figure S2**: Blue-white assays at 30°, with *rho*$^{+}$ *infB*$^{+}$ shelter plasmid pHYD5212, for *rho*-136$^{opal}$ *ruv* synthetic lethality and its suppression. General notations are as described in legend to Figure 1A; the designation *rho* refers to allele *rho*-136$^{opal}$. Experiments were on LB at 30°, with the exception of that for sub-panel viii which was on defined medium. UvsW and UvsW-K141R (for sub-panels iv and v, respectively) were each expressed from an IPTG-inducible $P_{tac}$ promoter on a chromosomally integrated construct (8). The medium for sub-panel v was supplemented with IPTG, whereas basal UvsW expression (without IPTG) was sufficient for the phenotype depicted in sub-panel iv; the apparently high proportion of white colonies in sub-panel iv is explained by the differential toxicity of UvsW in *rho*$^{+}$ derivatives, that is, in the blue colonies (8). Strains for the different sub-panels were (all strain numbers are prefixed with GJ): i, 15447; ii, 19853; iii, 19846; iv, 15465; v, 15466; vi, 15446; vii, 15490; and viii, 15485.

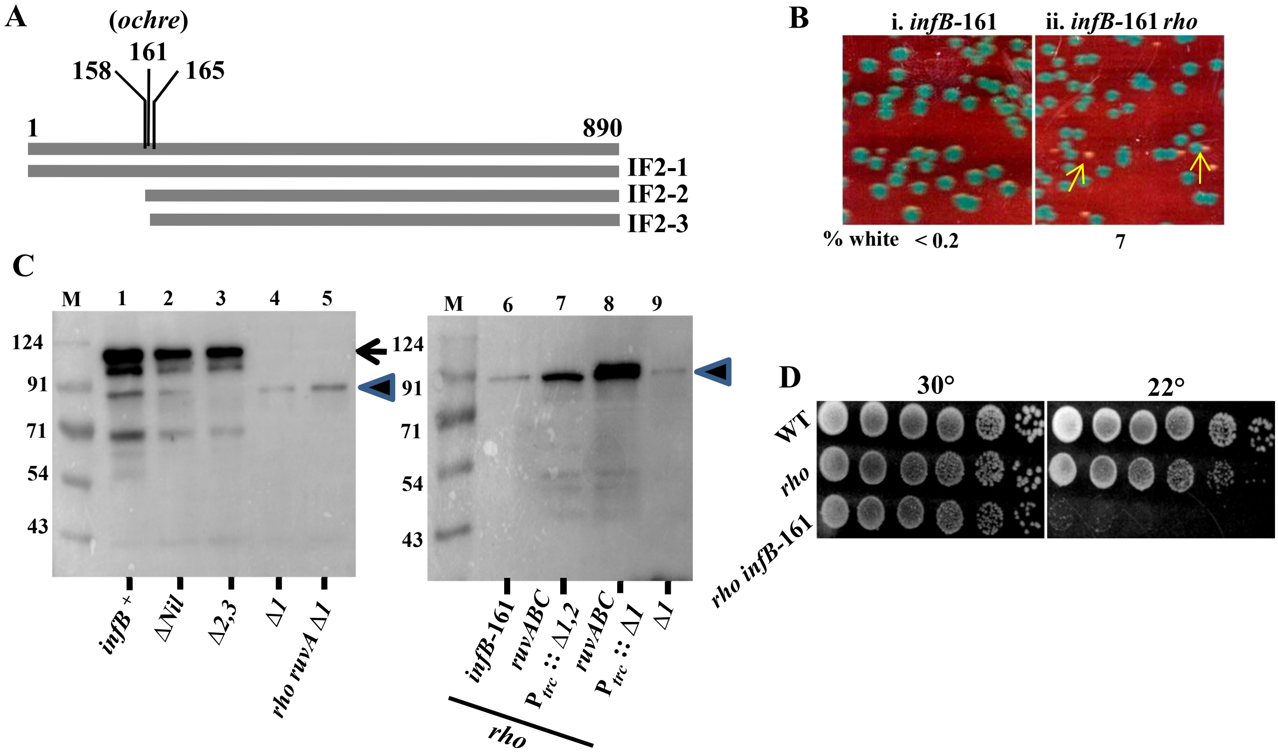


**Supplementary Figure S3**: IF2 isoforms and *infB*-161$^{ochre}$ mutation. Genotype notations *rho* and *infB*-161 in different panels, refer, respectively, to *rho*-136$^{opal}$ and *infB*-161$^{ochre}$. In all panels, strains whose designations include Δ*Nil*, Δ*1*, or Δ*2,3* were also Δ*infB*. All strain numbers mentioned are prefixed with GJ. **(A)** Linear representation of 890-codon-long *infB* open-reading frame; positions 1, 158, and 165 representing three in-frame initiation codons are marked. Depicted beneath are the three corresponding isoforms IF2-1, IF2-2, and IF2-3, respectively. Location of *ochre* mutation at position 161 is also marked. **(B)** Blue-white screening assay (with *rho*$^{+}$ *infB*$^{+}$ shelter plasmid pHYD5212) on LB at 30° of isogenic *rho*$^{+}$ (15449, left) and *rho*-136$^{opal}$ (15445, right) derivatives of *infB*-161$^{ochre}$ mutant. Other notations are as described in legend to Figure 1A. **(C)** Western blotting (following electrophoresis on 10% polyacrylamide gel with sodium dodecyl sulphate) with anti-IF2 antibody of strains whose genotypes/features are indicated beneath each lane. Positions of migration of IF2-1 and IF2-2,3 polypeptides are marked by arrow and arrowhead, respectively. M, lane for molecular mass standards, of sizes as shown at left (in kDa). Strains used for different lanes were: 1, 15410; 2, 19101; 3, 19102; 4, 15494; 5, 15499; 6, 15445; 7, 15458; 8, 15464; and 9, 15494. Cultures for lanes 7-8 were with IPTG supplementation. **(D)** Dilution spotting assay on defined medium at 30° and 22°. Strains for different rows were (from top, relevant genotypes at left; WT (wild-type): 15410, 15441, and 15445, respectively.

**Supp. Fig. S4**

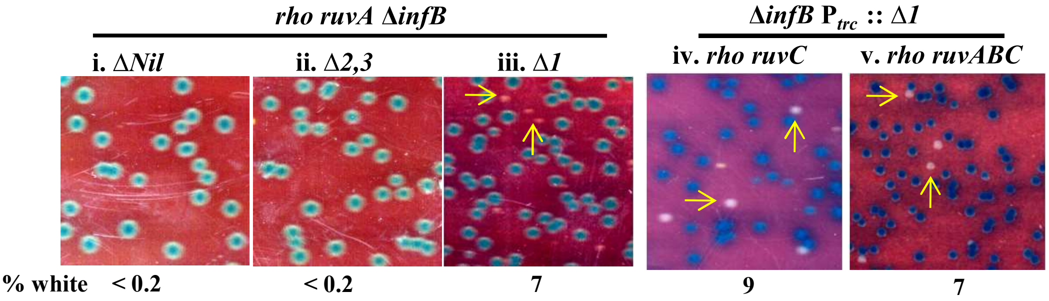


**Supplementary Figure S4**: Modulation of *rho ruv* synthetic lethality by differential expression of IF2 isoforms. Blue-white screening assays were performed at 30° (on LB and on defined medium for, respectively, sub-panels i-iii and iv-v), with strains carrying $rho^{+}$ $infB^{+}$ shelter plasmid pHYD5212. The medium for each of sub-panels iv and v was supplemented with IPTG. General notations are as described in legend to Figure 1A; the designation *rho* refers to allele *rho*-$136^{opal}$. Strains for the different sub-panels were (all strain numbers are prefixed with GJ): i, 15498; ii, 15500; iii, 15499; iv, 19152; and v, 15464.

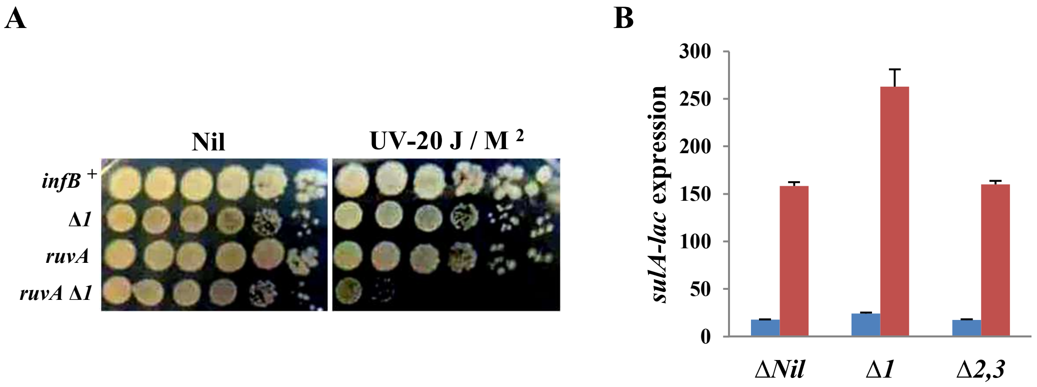


**Supplementary Figure S5**: UV tolerance and SOS induction in strains expressing different IF2 isoforms. Strains whose designations include Δ*Nil*, Δ*1*, or Δ*2,3* were also Δ*infB*. All strain numbers mentioned are prefixed with GJ. **(A)** Dilution-spotting assay on LB to determine UV-irradiation tolerance. Strains for different rows were (from top, relevant genotypes at left): 15410, 15494, 15488, and 19166, respectively. **(B)** *sulA-lac* expression values (mean and standard error in Miller units) as a measure of SOS induction in LB-grown cultures of the indicated strains. Blue and red bars represent cultures without and with phleomycin supplementation (at 3 µg/ml), respectively. Strains were: Δ*Nil*, 19839; Δ*1*, 19841; and Δ*2,3*, 19840.
